# Discovery of clinically approved drugs capable of inhibiting SARS-CoV-2 *in vitro* infection using a phenotypic screening strategy and network-analysis to predict their potential to treat covid-19

**DOI:** 10.1101/2020.07.09.196337

**Authors:** Douglas Ferreira Sales-Medina, Ludmila Rodrigues Pinto Ferreira, Lavínia M. D. Romera, Karolina Ribeiro Gonçalves, Rafael V. C. Guido, Gilles Courtemanche, Marcos S. Buckeridge, Édison L. Durigon, Carolina B. Moraes, Lucio H. Freitas-Junior

**Affiliations:** Department of Microbiology, Institute of Biomedical Sciences, University of Sao Paulo, Av. Prof. Lineu Prestes, 1374, Sao Paulo-SP 05508-900, Brazil; RNA Systems Biology Laboratory (RSBL), Post-Graduate Program in Cell Biology, Institute of Biological Sciences, Federal University of Minas Gerais & INCT-Vacinas, Av. Antônio Carlos, 6627, Belo Horizonte-MG 31270910, Brazil; São Carlos Institute of Physics, University of Sao Paulo, Avenida João Dagnone, 1100, São Carlos, SP 13563-120, Brasil São Carlos, Brazil; Bioaster, 28, rue du Docteur Roux, 75015 Paris, France; Department of Botany, Institute of Biological Sciences, University of Sao Paulo, Rua do Matão, 277, Sao Paulo-SP 05508-900, Brazil; Department of Pharmaceutical Sciences, Federal University of Sao Paulo, Rua São Nicolau, 210, Diadema – SP 09913-030, Brazil

**Keywords:** SARS-CoV-2, covid-19, drug repurposing, antivirals, phenotypic screening, network analysis

## Abstract

The disease caused by SARS-CoV2, covid-19, rapidly spreads worldwide, causing the greatest threat to global public health in the last 100 years. This scenario has become catastrophic as there are no approved vaccines to prevent the disease, and the main measures to contain the virus transmission are confinement and social distancing. One priority strategy is based on drug repurposing by pursuing antiviral chemotherapy that can control transmission and prevent complications associated with covid-19. With this aim, we performed a high content screening assay for the discovery of anti-SARS-CoV-2 compounds. From the 65 screened compounds, we have found four drugs capable to selectively inhibit SARS-CoV-2 *in vitro* infection: brequinar, abiraterone acetate, neomycin, and the extract of *Hedera helix*. Brequinar and abiraterone acetate had higher inhibition potency against SARS-CoV-2 than neomycin and *Hedera helix* extract, respectively. Drugs with reported antiviral activity and in clinical trials for covid-19, chloroquine, ivermectin, and nitazoxanide, were also included in the screening, and the last two were found to be non-selective. We used a data mining approach to build drug-host molecules-biological function-disease networks to show in a holistic way how each compound is interconnected with host node molecules and virus infection, replication, inflammatory response, and cell apoptosis. In summary, the present manuscript identified four drugs with active inhibition effect on SARS-CoV-2 *in vitro* infection, and by network analysis, we provided new insights and starting points for the clinical evaluation and repurposing process to treat SARS-CoV-2 infection.

**Summary sentence:** Discovery of drug repurposing candidates, inhibitors of SARS-CoV-2 infection *in vitro*, using a phenotypic screening strategy and network analysis.

## Introduction

The world is experiencing an unprecedented pandemic of a novel severe acute respiratory syndrome coronavirus 2 (SARS-CoV-2). The first infected patients were reported in Wuhan, Hubei province of China, in December 2019. The disease caused by SARS-CoV-2, covid-19 ^(1,2)^ has high mortality and morbidity and rapidly spread, creating an enormous impact on the health systems worldwide. As of June 2020, over 8 million cases of SARS-CoV-2 infection were confirmed with at least 450 thousand deaths officially recorded worldwide ^(3)^. SARS-CoV-2 is an enveloped, positive-sense singlestranded RNA betacoronavirus of the family *Coronaviridae*. Infection with most members of this family cause only mild, common cold-like symptoms. SARS-CoV-2 is part of the family the first SARS-CoV, which caused in 2002 approximately 8000 cases, with a fatality rate of 10%, and the Middle East respiratory syndrome coronavirus (MERS-CoV), which produced in 2012 around 2500 confirmed cases with a fatality rate of 36% ^(4,5,6)^. Compared to MERS or SARS, SARS-CoV-2 appears to spread more efficiently, making it difficult to contain and increasing its pandemic potential ^(6,7,8)^.

While the discovery and development of new antivirals that can efficiently treat covid-19 should be actively pursued, drug repositioning stands out as an attractive strategy to be explored for SARS-CoV-2. Toward this end, many preclinical studies and investigational clinical trials have been conducted worldwide to determine which therapeutic options readily available can serve not only as an antiviral therapy but also as a cure of covid-19, including drugs that can efficiently revert the inflammatory condition that develops in association with severe cases, as recently reported for dexamethasone ^(9)^.

Some screening campaigns of approved drug libraries and bioactive molecules have been conducted for the discovery of SARS-CoV-2 inhibitors. Antiparasitic drugs as chloroquine and its derivative hydroxychloroquine ^(10,11)^, ivermectin ^(12)^, nitazoxanide ^(10)^ have shown antiviral activity against SARS-CoV-2 *in vitro*. Another approved drug that decreased the coronavirus infection in cell culture was azithromycin ^(13)^, a widely used broad-spectrum antibiotic. Nucleoside analogs as remdesivir, favipiravir, arbidol, ribavirin have already been tested for activity against SARS-CoV-2 ^(10, 14–16)^. The antiviral lopinavir, a protease inhibitor, has also been shown inhibitory activity of SARS-CoV-2 infection in Vero E6 cells ^(14)^.

Over the years, data mining based-computational analyses have also been used as hypothesis-generating research helping to fasten the repurposing approach by redirecting the *in vitro* and *in vivo* testing and chosen potential drugs that could lead to significant results and find successful treatments for different diseases ^(17)^. By building a drug-molecules-disease/function network we show in this study in a holistic way how the drugs tested in vitro are interconnected with node molecules and biological functions associated with *Coronaviridae* replication. These findings provide insights toward possible mechanisms of action of these drugs and may point to target molecules as alternative therapeutic options to SAR-COV-2.

Given the current covid-19 pandemic scenario, there is an urgent need to find therapeutic solutions that limit viral infection and reduce clinical complications. We have developed a high content screening assay to identify inhibitors of SARS-CoV-2 infection in vitro and used to screen 65 compounds, some of them previously reported as having broadspectrum antiviral activity. Computational analysis was used to build drug-host molecules-biological function-disease networks based on the Ingenuity Knowledge Base (IKB), a manually and automatically curated and extracted repository of biological interactions and functional annotations created from millions of individually modeled relationships between proteins, genes, complexes, cells, tissues, drugs and disease extracted from the literature.

## Materials and Methods

### Virus and cell lines

The SARS-CoV-2 used in this study (HIAE-02: SARS-CoV-2/SP02/human/2020/BRA, GenBank Accession No. MT126808.1) was isolated from a nasopharyngeal sample of a confirmed covid-19 patient in the Hospital Israelita Albert Einstein, São Paulo, Brazil. The virus was passaged twice in Vero E6 cell line (ATCC) maintained in DMEM High Glucose (Sigma-Aldrich) supplemented with 2% heat-inactivated Fetal Bovine Serum (Thermo Scientific) and 100 U/mL of penicillin and 100 μg/mL of Streptomycin (Thermo Scientific). Cells were maintained at 37 °C with 5% CO_2_. The supernatant of infected cells was stored in aliquots at - 80 °C, and the viral titer was determined by plaque assay in Vero CCL-81. Briefly, 1×10^5^ Vero CCL-81 cells were seeded on each well of a 24-well plate in DMEM High Glucose supplemented as described above at 37° C with 5% CO_2_. After 24 h, the medium was removed, and 400 μL of medium with serial dilutions of SARS-CoV-2 were incubated for 1 h at 37 °C with 5% CO_2_ for virus adsorption. Then, the medium was removed and replaced with 500 μL of DMEM High Glucose supplemented as above added 3% of carboxymethyl cellulose and incubated for 72 h. After that, the medium was removed, the plate was fixed with 4% paraformaldehyde in PBS (m/v) pH 7.4 for 15 min and stained with crystal violet 1% in 10% ethanol (m/v/v). The number of plaques was visually assessed, counted, and virus titer was calculated as plaque-forming units (PFU)/mL. All procedures involving the SARS-CoV-2 virus were performed in the biosafety level 3 laboratory at the Institute of Biomedical Sciences of the University of São Paulo.

### Compounds

Brequinar sodium salt (CAS No. 96201-88-6), Chloroquine diphosphate salt (CAS No. 50-63-5), U-73343 (CAS No. 142878-12-4), 6-Azauridine (CAS No. 54-25-1), 5-Fluorouracil (CAS No. 51-21-8), Benztropine mesylate (CAS No. 132-17-2) and Bafilomycin A1 (CAS No. 88899-55-2) were purchased from Sigma-Aldrich. Active pharmaceutical ingredients for an additional 59 drugs, listed in Table S1, were provided by Eurofarma Laboratories, Brazil. Compounds were resuspended in Dimethyl sulfoxide (DMSO) (Sigma-Aldrich) for a final concentration of 10 mM or 10 mg/mL (for *Hedera helix* extract) and stored at - 20 °C. For the preparation of screening plates, compounds were manually pipetted on a 384-well polypropylene plate (Greiner Bio-One) and serially diluted in neat DMSO by a factor of two, starting from 10 mM or 10 mg/mL, and frozen at - 20 °C until use.

### Phenotypic screening assay

An amount of 2000 Vero E6 cells were seeded on each well of a 384-well assay plate (Greiner Bio-One) in 40 μL of DMEM High Glucose (Sigma-Aldrich) supplemented with 10% heat-inactivated Fetal Bovine Serum (Thermo Scientific), 100 U/mL of penicillin and 100 μg/mL of streptomycin (Thermo Scientific) at 37 °C, 5% CO_2_ for 24 h. After this period, the medium was aspirated using Biotek Multiflo FX, and 30 μL of DMEM High Glucose (Sigma-Aldrich) was added in each well. Serially-diluted compounds were manually transferred into a polypropylene 384-well plate (Greiner Bio-One) containing sterile phosphate-buffered saline (PBS) pH 7.4, for a final dilution factor of 33,3. Then, 10 μL of each well in the compounds plate was transferred to the cell-containing assay plate, followed by the addition of SARS-CoV-2 viral particles to the cells at the multiplicity of infection (MOI) 0.01 in 10 μL of DMEM High Glucose per well. DMSO-treated infected cells and DMSO-treated non-infected cells were used as controls. The assay plate was incubated for 1 h at 37 °C, 5% CO_2_ for virus adsorption, followed by the addition of 10 μL of DMEM High Glucose supplemented with 12% Fetal Bovine Serum per well. Thus, final concentrations in the assay plate were 0.5% DMSO and 2% FBS. After 48 h of incubation, the plates were fixed in 4% paraformaldehyde in PBS pH 7.4 and subjected to indirect immunofluorescence detection of viral cellular infection. After washing twice with PBS pH 7.4, plates were blocked with 5% bovine serum albumin (BSA) (Sigma-Aldrich) in PBS (BSA-PBS) for 30 min at RT and washed twice with PBS. As a primary antibody, either serum from a convalescent covid-19 Brazilian patient diluted 1:500 in PBS or a polyclonal rabbit antibody anti-SARS-CoV-2 nucleocapside protein (GeneTex) at 2 μg/mL in PBS were used to detect SARS-CoV-2 infection in Vero cells. The primary antibodies were incubated for 30 min, and plates were washed twice with PBS. As secondary antibodies, goat anti-human IgG labeled with FITC (Chemicon) or goat anti-rabbit IgG labeled with Alexa 488 (Thermo Scientific) was used diluted at 4 μg/mL in PBS and incubated for 30 min with 5 μg/mL 4’,6-Diamidine-2’-phenylindole dihydrochloride (DAPI, Sigma-Aldrich) in PBS to stain nuclei. The plates were washed twice with PBS and imaged in the Operetta High Content Imaging System (Perkin Elmer) using a 20x magnification objective. Four images were acquired per well.

### Data analysis

Acquired images were analyzed in the software Harmony (Perkin Elmer), version 3.5.2. Image analysis consisted of identifying and counting Vero E6 cells based on nuclear segmentation and viral infection based on the cytoplasmic staining detected by the immunofluorescence assay. The infection ratio (IR) was calculated as the ratio between the number of infected cells and the number of total cells counted in each well. The cell survival rate was calculated as the number of cells counted in each well divided by the average number of cells in the positive control (DMSO-treated infected cells) wells, multiplied by 100. The antiviral activity was determined by the normalization of the IR to the negative control (DMSO-treated infected and non-infected cells), as described. Concentration-response curves were plotted using the normalized activity and cell survival of each concentration. These two parameters were used to calculate the concentration of EC_50_ and CC_50_, compounds concentrations that reduce the infection ratio, and cell survival in 50%, respectively, compared to non-treated infected controls of each compound using GraphPad Prism version 7.0 (GraphPad Software, USA).

### Functional and network analysis

Functional and network analysis was performed with Ingenuity Pathway Analysis (IPA, Qiagen). The network was built considering only direct and indirect experimentally validated node relationships available from the database and direct acquisition from the literature by Ingenuity Knowledge Base (IKB). Fisher’s exact test measured the significance of the association between drug-node-function and disease. As a result, a P-value is obtained, determining the probability that the association between the nodes in the networks generated can be explained by chance alone. The analysis was performed based on IKB content of date 2020–06. The Molecule Activity Predictor (MAP) tool from IPA was used to predict the upstream and/or downstream effects of activation or inhibition of molecules in a network or pathway given one or more neighboring molecules with “known” activity from the literature. MAP enabled us to visualize the overall effect on a pathway or network and, therefore, to generate a more accurate hypothesis.

The molecules or functions that are predicted to be activated are colored in orange. Those predicted to be inhibited (colored blue), are based on their interactions with molecules known (or manually assigned by the user) to be activated or inhibited. The predictions become less defined as the distance from the node(s) with known activation state increases represented by tones of orange and blue (darker to fainter) from the drug, expressed in red.

## Results

### High Content Screening Assay Development

A high content screening (HCS) assay was developed to measure the effects of 65 drugs and experimental compounds on infection and cytotoxicity *in vitro* on Vero E6 cells infected with a Brazilian isolate of SARS-CoV-2, aiming at providing candidates for clinical repurposing and chemical probes for validation of drug targets for covid-19. Figure 1A shows a graphical representation of the overall workflow of the assay’s conditions and procedures to screen the drug library. Compounds were evaluated in concentration-response using HCS data and classified for their antiviral activity into non-active, non-selective/active, and selective/active. Computational analysis was used to build a drug-host molecule-biological function-disease network for selective drugs with novel anti-SARS-CoV-2 activity. Figures 1B (a to d) show representative images from the automated high content image analysis used to quantify cell infection performed in optimized assay conditions, with nuclei and cytoplasm segmentation, followed by classification of cells into infected (green) and non-infected (red). The immunofluorescence detection of SARS-CoV-2 intracellular was performed using the serum from a convalescent covid-19 patient that proved to have higher sensitivity than a commercial anti-SARS-CoV-2 N protein polyclonal rabbit antibody (Figure S1).

**Figure 1.**
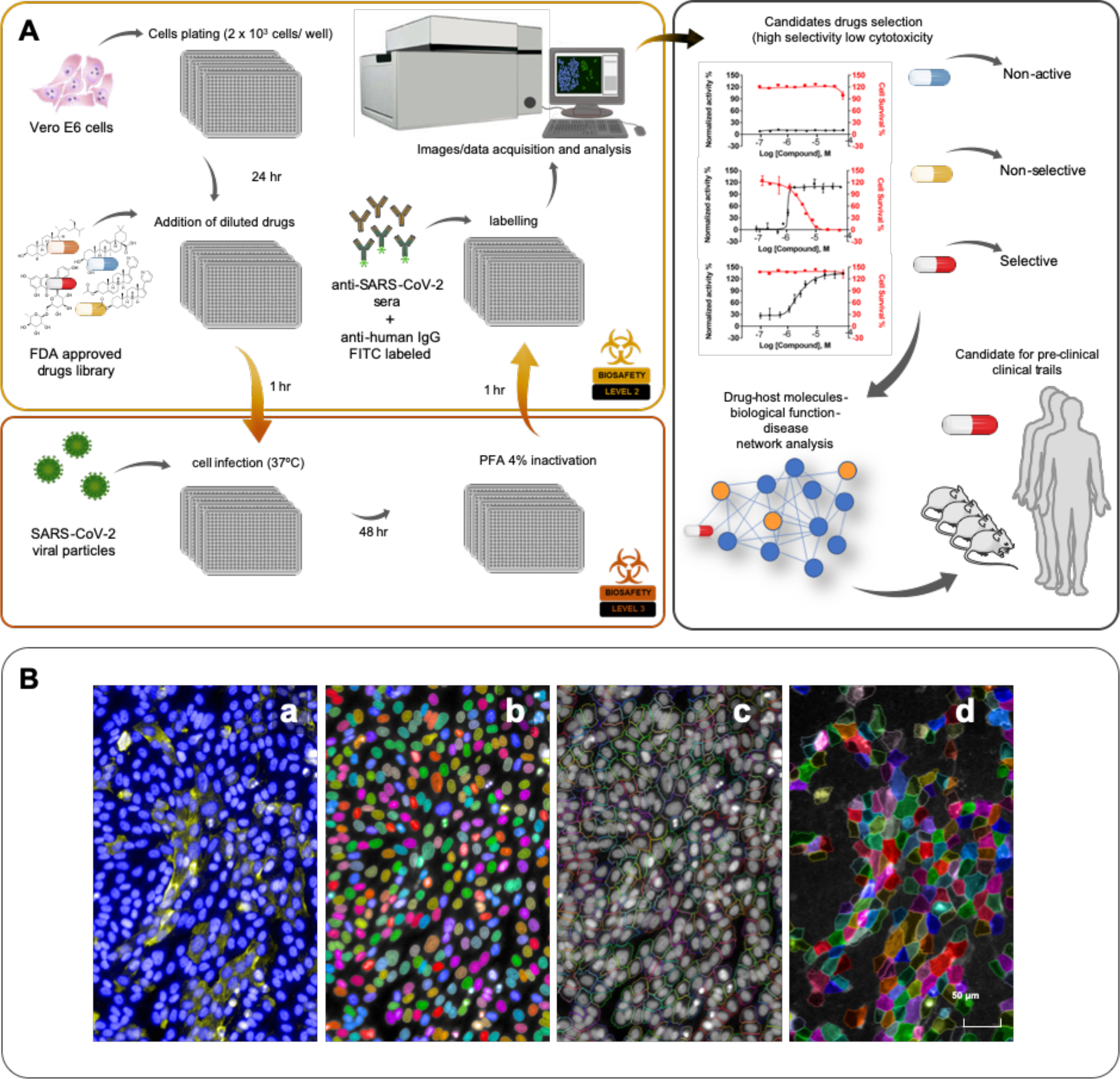
High Content Screening for SARS-CoV-2 antiviral discovery. **(A)** Top left and right panels: High Content Screening Assay (HCSA) experiment workflow graphical representation for the identification of drugs with selective activity against SARS-CoV-2 *in vitro* infection in Vero E6 cells. A network-based analysis is performed to predict the biological interactions and downstream effects of each selected lead drug/compound on host node molecules for drug repurposing in the context of viral infections; **(B)** Representative images from the HCSA immunofluorescence-based assay of SARS-CoV-2 *in vitro* infection of Vero cells infection showing: **(a)** DAPI stained (blue) cell nuclei with SARS-CoV-2 infection cytoplasmic localization (yellow); **(b)** Automated nuclei and **(c)** cytoplasm segmentation localization based on their respective staining; **(d)** Automated identification and selection of viral infection based on detection of cytoplasmic immunofluorescence staining of viral particles.

### Evaluation of drugs with known anti-SARS-CoV-2 activity

The assay was validated with three antiparasitic drugs with previously reported antiviral activity against SARS-CoV-2 *in vitro* infection for covid-19: chloroquine ^(10)^, ivermectin ^(12)^ and nitazoxanide ^(18)^. All three drugs were capable of inhibiting SARS-CoV-2 infection in a concentration-dependent manner (Figure 2), with EC_50_s of similar range (4.7 μM for chloroquine, 1.0 μM for nitazoxanide and 1.7 μM for ivermectin – Table 1). However, the evaluated CC_50_ for nitazoxanide and ivermectin were at the same range (3.3 and 2.2 μM, respectively), suggesting that they are not selective. Only chloroquine was non-cytotoxic, with a selectivity index (SI) greater than 18, while both nitazoxanide and especially ivermectin displayed cytotoxicity, with poor SI values (3.3 for nitazoxanide and 1.3 for ivermectin). The poor selectivity of these two drugs *in vitro* is further evidenced by the increasing frequency of apparent pyknotic nuclei at higher drug concentrations (red arrows in Fig. 2) and reduced cell number, as opposed to reduced incidence of host cell diving nuclei (green arrows in Fig. 2).

**Figure 2.**
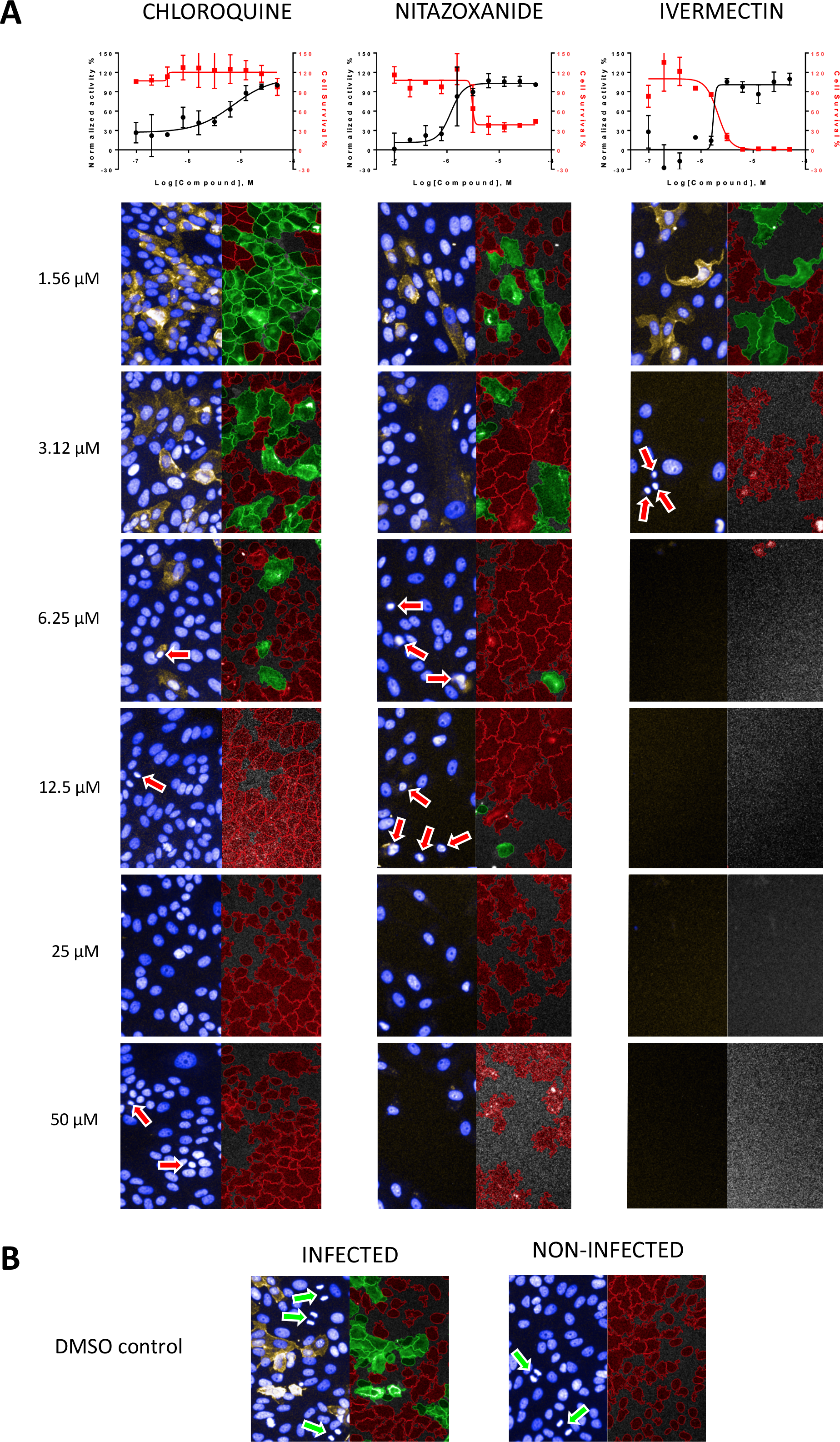
Comparison of different concentrations of chloroquine, ivermectin, and nitazoxanide by HCS. (A) Representative images of different concentrations of chloroquine, ivermectin, and nitazoxanide captured with a 20x objective. On the top of the image, the graphs represent each compound activity, followed by the images representing each concentration in a crescent manner. The images on the left it is possible to identify nuclei (stained with DAPI – blue) and the virus (represented in yellow); the images on the right represents the analysis of software Harmony, where in red are uninfected cells and in green the infected ones. Arrows on redpoint to apoptotic nuclei. (B) Representative images of infected and non-infected controls, with the same stain as described above. Green arrows point to cells in mitosis.

**Table 1:** Activity of antiviral compounds against SARS-CoV-2 infection *in vitro*.

|  | **EC<sub>50</sub>, μM** | **CC<sub>50</sub>, μM** | **SI** |
| --- | --- | --- | --- |
| Chloroquine | 4.7 | > 50 | 18.3 |
| Nitazoxanide | 1.0 | 3.3 | 3.3 |
| Ivermectin | 1.7 | 2.2 | 1.3 |
| Brequinar | 0.3 | > 50 | 166.7 |
| U-73343 | 2.7 | 9.6 | 3.6 |
| 6-Azauridine | > 50 | > 50 | > 50 |
| 5-Fluorouracil | > 50 | > 50 | ND |
| Benztropine | > 50 | > 50 | ND |
| Bafilomycin A1 | > 50 | > 50 | > 50 |

SI, selectivity index

Other experimental compounds and drugs with known antiviral activity, but not previously reported for SARS-CoV-2, were also tested in the HCS assay. Among these, only brequinar ^(19, 20, 21)^ and the experimental compound U-73343 ^(22)^ showed activity against SARS-CoV-2 *in vitro*, with EC_50_s of 0.3 μM for brequinar and 2.7 μM for U-73343. However, only brequinar was sufficiently selective, with SI higher than 166, while U-73343 had a SI of 3.6. The other compounds, 6-Azauridine ^(23, 24)^, 5-fluorouracil ^(23, 25)^, benztropine ^(26)^, bafilomycin A1 ^(27, 28)^, showed no activity against SARS-CoV-2 (Table 1).

### Screening of drugs for repurposing for covid-19

The assay was then used to screen the other 56 drugs currently used in the clinic for several indications. Among these, two drugs showed new activity against SARS-CoV-2 (Table 2): the antiandrogen abiraterone acetate (EC_50_ of 7.1 μM) and the aminoglycosidic antibiotic neomycin (EC_50_ of 12.9 μM). Besides these two molecules, moderate antiviral activity against SARS-CoV-2 was also observed for the extract of *Hedera helix* leaves, used as a treatment for cough, with an EC_50_ of 51 μg/mL. These compounds exhibited no cytotoxic effect under the assay conditions, thereby being sufficiently selective against SARS-CoV-2. Other tested compounds showed no activity against SARS-CoV-2 *in vitro* (Table S1).

**Table 2:** Activity of new drugs against SARS-CoV-2 infection *in vitro*.

|  | **EC<sub>50</sub>** | **CC<sub>50</sub>** | **SI** |
| --- | --- | --- | --- |
| Abiraterone acetate | 7.1 μM | > 50 μM | > 7.0 |
| Neomycin | 12.9 μM | > 50 μM | > 3.9 |
| *Hedera helix* | 51 μg/mL | > 50 μg/mL | > 1.0 |

SI, selectivity index

Figure 3 shows the concentration-response curves for antiviral and cytotoxic activity of the most selective compounds: brequinar, abiraterone, *Hedera helix*, and neomycin, along with representative images of intracellular infection under active and non-active drug concentrations.

**Figure 3.**
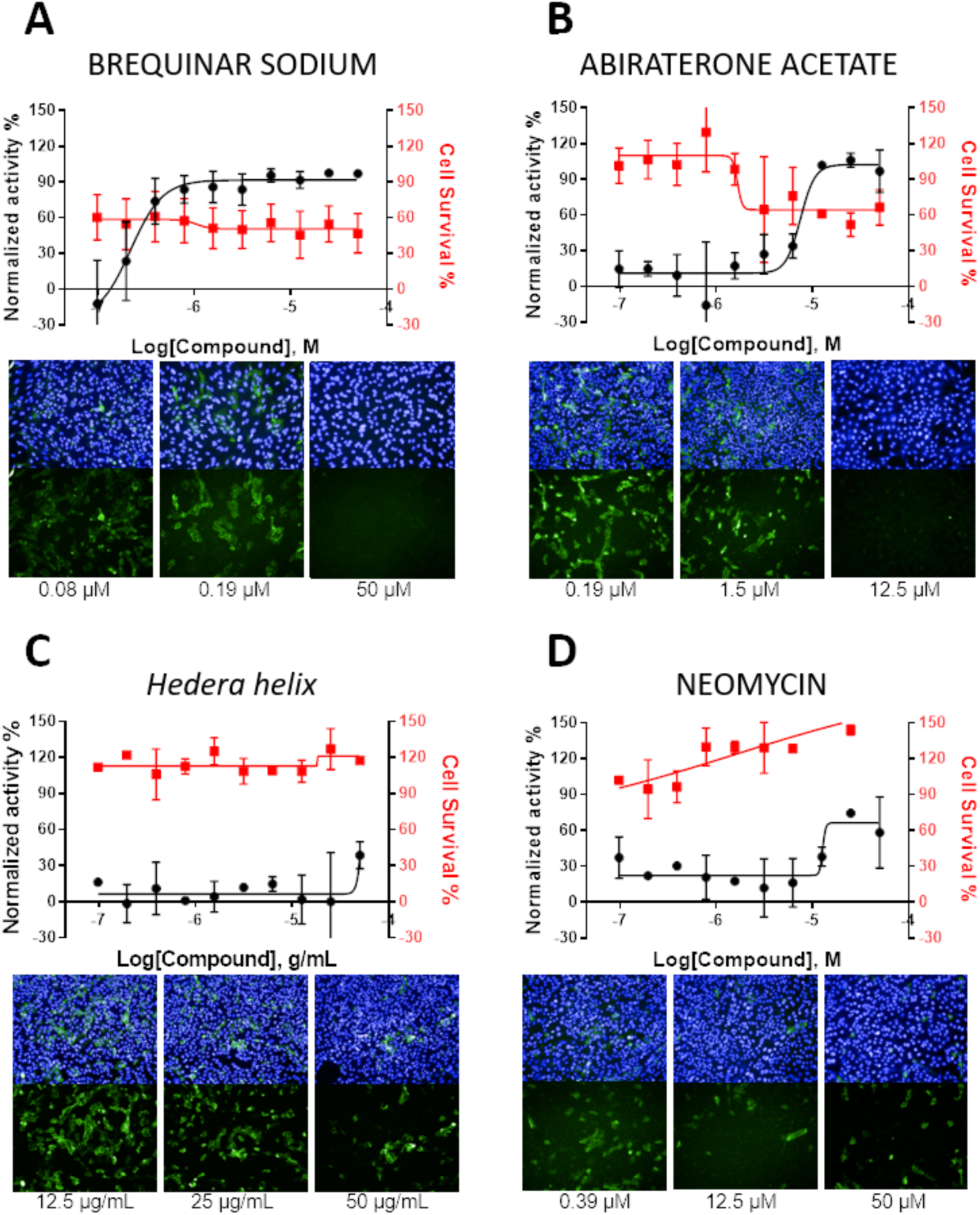
Antiviral and cytotoxic activities of compounds against SARS-CoV-2. Antiviral activity is shown in the black curves, and cytotoxic activity is shown in red curves. Data points are mean values, and bars are the standard deviation of two independent experiments. Images of infection in the presence of different compound concentrations for each compound, with SARS-CoV-2 stained in green (shown in upper and lower panels) and cell nuclei in blue (shown only in upper panels).

### Network prediction analysis of drug-host molecules-biological function-diseases interaction

A data mining approach was used to build a drug/compound-host molecules interaction network, as shown in Figure 4 abiraterone acetate, neomycin, brequinar and some of the main components found in the *Hedera helix* extract, e.g. hexa-D-arginine, rutin, quercetin, hederagenin, nicotiflorin, kaempferol, stigmasterol, and alpha-hederin. All networks built have the same number of nodes that represent host molecules, biological functions (“pro-inflammatory cytokines,” “cell apoptosis) and disease (viral infection and RNA virus replication) without counting those related to the drug/compound. The nodes are listed in Table S2. The networks were built by using an IPA tool called “Grow,” where new node molecules are added to the growing network in order of their interconnectedness and also specific connectivity based on their position in a more extensive network stored on IPA database (Global Molecular Network – GMN – gene that is most connected to the growing network). The nodes in the networks are interconnected directly and/or indirectly represented as solid and dashed lines (edges), respectively. We show networks built by growing from the functions and disease nodes and manually adding the drug/compound. When the drug is present in the network, its node is colored in red. The prediction analysis revealed the potential role/connection of each one of the drugs with other nodes/molecules. When nodes and/or biological functions/diseases are colored in tones of orange, they are predicted to be downstream activated and inhibited when colored in blue tones. Importantly, the network considers node molecules with more than 10 independent connections with others, suggesting that those molecules might be central in coronavirus replication. The nearest neighbors of each drug/compound node(s) and their edges are in bold. Our predictions show that the drugs abiraterone acetate (Figure 4A) and the compounds present in the extract of *Hedera helix* (Figure 4D) have a potential downstream inhibitory activity in all nodes related to biological functions and diseases (colored in blue). Alternatively, for the drugs brequinar (Figure 4B) and neomycin (Figure 4C), the nodes related to cell apoptosis and pro-inflammatory cytokines production are predicted to be activated (colored in orange), respectively. The nearest neighbors (molecules) of each drug/compound node connected directly or indirectly are called focus molecules. For the drug abiraterone acetate (Figure 4A), we found four focus molecules: the enzymes cytochrome P450 family 17, subfamily A member 1 (CYP17A1), transmembrane serine protease 2 (TMPRSS2), kallikrein-related peptidase 3 (KLK3) and the androgen receptor (AR). All four nodes are colored in blue, based on literature findings that show that the drug abiraterone has an inhibitory effect by decreasing their activation and or expression. For neomycin (Figure 4B) we found eleven focus molecules: the caspases 8 and 9 (CASP8, CASP9), CXC-motif chemokines 10 and 11 (CXCL10, CXCL11), interleukin-10, Src family tyrosine kinase (LCK), the mitogen-activated protein kinases 1 and 3 (MAPK1, MAKPK3), phosphatidylinositol (PI) and reactive oxygen species (ROS), showing that this drug has a pro-inflammatory activity. Brequinar has three focus genes; two predicted to be activated, e.g., integrin subunit alpha M (ITGAM) and Fc-gamma receptor 1 (FcγRI) and one inhibited, the molecular target of brequinar, the enzyme dihydroorotate dehydrogenase (DHODH). For the *Hedera helix* extract, the network was built with its eight major components. The 29 focus genes interconnected with *Hedera helix* compounds are listed in Table S2. Nevertheless, we could identify a central node that is interconnected with a significant number of other nodes, the molecule Furin, which is predicted to be inhibited. Three compounds from the *Hedera helix* extract have inhibition activity in this basic-amino-acid-specific peptidase, e.g., hexa-D-arginine, rutin, and quercetin. Also, *Hedera helix* compounds showed anti-inflammatory activity, as all the cytokines in the network are predicted to be inhibited, e.g., Tumor Necrosis Factor Alpha (TNF), interleukin 1 beta (IL1B), interleukin 6 (IL6).

**Figure 4-.**
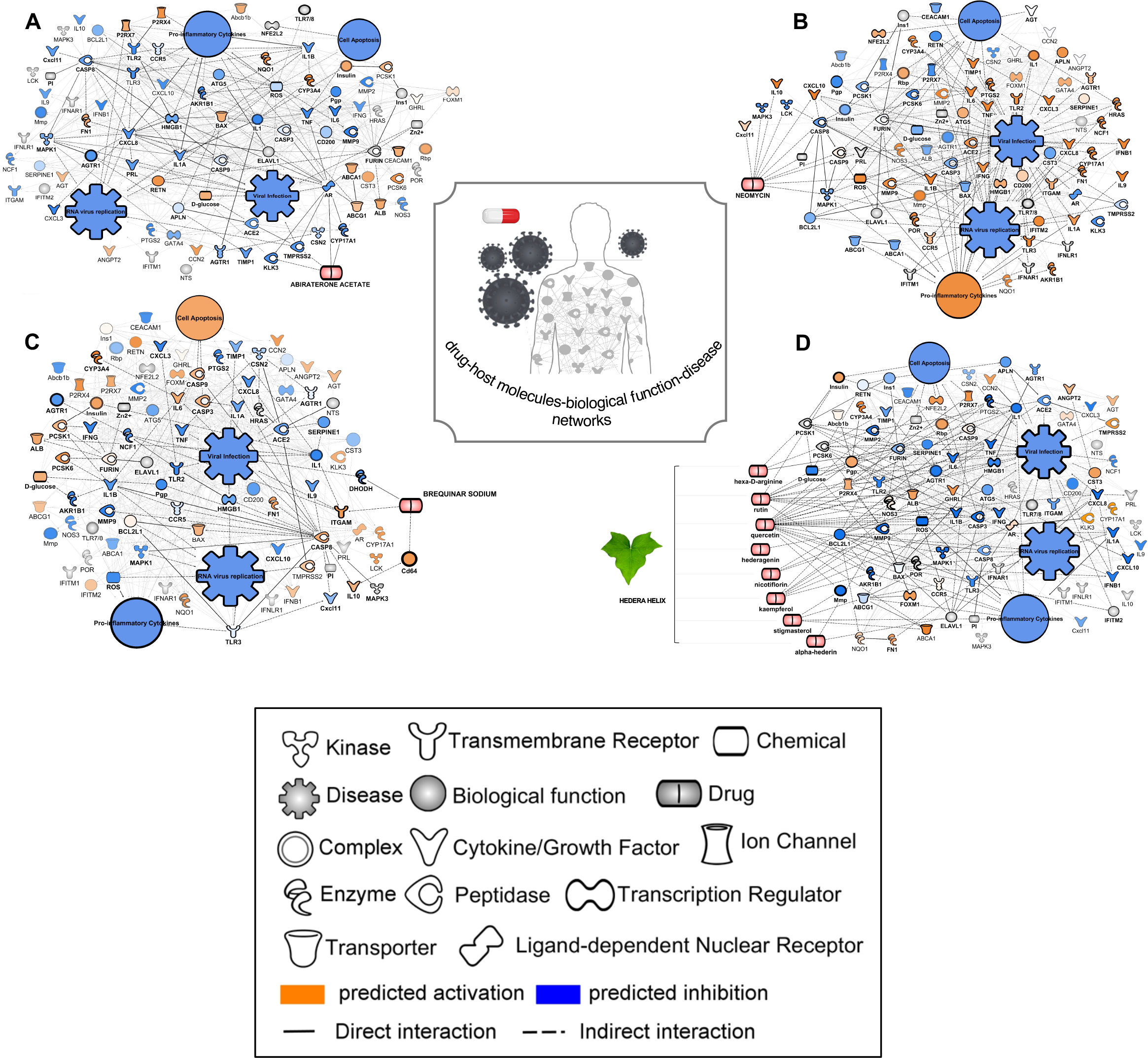
Drug/compound-host molecules-biological function-disease-networks. Networks built with the drugs abiraterone acetate (**A**), neomycin (**B**), brequinar sodium (**C**), and the main compounds found in the Hedera helix extract e.g. hexa-D-arginine, rutin, quercetin, hederagenin, nicotiflorin, kaempferol, stigmasterol and alpha-hederin (**D**). The presence of these drug/compounds has a potential inhibition or activation effect on the network molecules, colored in blue and orange, respectively. The gray-colored nodes represent molecules without the predicted activation state. Nodes have different shapes representing their different classification, type.

## Discussion

The urgent search for treatments for covid-19 has hastened the repurposing drugs process and initiation of several clinical tests with drugs with reported *in vitro* antiviral activity for SARS-CoV-2 and instigated the approval of expanded access and compassionate use of these drugs to treat critically ill patients, even if they have not yet gone through all the steps necessary for specific use for covid-19.

In this report, we describe the use of a cell-based phenotypic screening assay to test the activity of drugs/compounds against SARS-CoV-2 *in vitro* infection with known antiviral activity or approved for human use. Our assays also included drugs previously tested against SARS-CoV-2, as a proof of test, like the drug chloroquine, widely used as antimalarial agents worldwide and described in screenings of FDA-approved drugs as active agents against different viral infections and recently for SARS-CoV-2. Here we report *in vitro* activity and propose chemotherapy candidates for covid-19. Chloroquine (and its derivative hydroxychloroquine) as an anti-SARS-CoV-2 treatment is still a topic with much controversy and debate ^(29–34)^. Other two antiparasitic drugs, ivermectin and nitazoxanide have been previously described as having antiviral activity, including for SARS-CoV-2. Nitazoxanide has been commercialized in Latin America and India for the treatment of a broad spectrum of intestinal infections. In recent years, nitazoxanide and its analogs have been proposed as a new class of antiviral agents. Tizoxanide, a nitazoxanide metabolite, can inhibit the maturation of rotavirus viral protein 7 (VPN) ^(35)^ and the replication of flaviviruses, including dengue serotype 2, Japanese encephalitis, yellow fever viruses ^(36, 37)^, influenza virus and rotavirus ^(35)^. Nitazoxanide has antiviral activity against MERS-CoV ^(38)^ and was able to block SARS-CoV-2 *in vitro* infection at low-micromolar concentrations ^(10)^. It also has been shown to induce type I IFN pathways by enhancing the RNA sensing axis that is usually trigged by foreign RNA cytoplasmic exposure ^(39)^. Ivermectin has been described as a SARS-CoV-2 inhibitor with *in vitro* activity ^(40, 41)^. This broad-spectrum parasiticide has exhibited antiviral activity against several RNA viruses, such as influenza A, zika, West Nile, chikungunya, yellow fever, and dengue ^(42, 43)^. The proposed mechanism of action for ivermectin is to inhibit viral replication by specifically targeting the activity of non-structural 3 helicase (NS3 helicase) *in vitro* ^(44)^. The use of ivermectin to treat covid-19 patients was evaluated in an observational registry-based study involving critically ill SARS-CoV-2-infected patients. The treatment was found to be associated with a lower mortality rate and reduced healthcare resources use ^(41, 45)^. We confirm here that both ivermectin and nitazoxanide have antiviral activity *in vitro* against SARS-CoV-2, but both were non-selective under these experimental conditions.

Brequinar was identified as the most potent compound with inhibitory activity against SARS-CoV-2 *in vitro*, with EC_50_ in the submicromolar range. Brequinar was first described as having antineoplastic properties because of its cytostatic effects on rapidly dividing cells and also enhance the *in vivo* antitumor effect of other antineoplastic agents ^(44)^. Here we show in the brequinar network, the nodes related to viral infection as well as the pro-inflammatory cytokines are predicted to be inhibited. The node related to cell apoptosis in predicted to be activated, colored in orange. We also identified by network analysis that brequinar is directly connected with the node for the enzyme dihydroorotate dehydrogenase (DHODH), an enzyme responsible for de novo pyrimidine biosynthesis. Brequinar was described as an inhibitor of DHODH, thereby blocking *de novo* biosynthesis of pyrimidines ^(46, 47)^, nucleosides necessary for replication of the viral genome ^(48–51)^. In another phenotypic assay conducted under similar conditions as the one reported herein, we verified that brequinar was active *in vitro* against the yellow fever virus (EC_50_ of 10.5 μM), albeit with lower potency than determined for SARS-CoV-2 herein ^(22)^. These findings suggest that SARS-CoV-2 might be more dependent on pyrimidines (uridine and cytosine) for its replication, which is in line with the fact that the SARS-CoV-2 virus has a high uracil content ^(52)^. The DHODH inhibition might slow the SARS-CoV-2 RNA polymerase activity due to the host UTP depletion, thereby affecting viral RNA replication. Luthra et al. 2018 ^(53)^ also demonstrated the brequinar capacity of blocking *in vitro* infection by the Ebola virus (EBOV), vesicular stomatitis virus (VSV), and Zika (ZIKV). The same work demonstrated that brequinar also exerts antiviral activity through the induction of expression of interferon-stimulated genes (ISGs), which encode for a variety of antiviral effectors. Hadjadj et al. 2020 ^(54)^ have shown that critically ill covid-19 patients have profoundly impaired type I IFN response, characterized by low IFN production and activity. Thus, it can be hypothesized that brequinar could also be a strong candidate to treat covid-19 as it could potentially induce an antiviral response by inducing ISG expression and restore type I interferon levels.

Abiraterone acetate also showed anti-SARS-CoV-2 activity in our study, confirming recent previous results from a study that has also demonstrated its activity against SARS-CoV-2 *in vitro* infection ^(55)^. Our network analysis showed the potential downstream effects of the drug abiraterone acetate, and biological function nodes, as well as those related to viral infection, are predicted to be inhibited by abiraterone acetate. The nodes directly connected to abiraterone acetate are the enzymes CYP17A1, TMPRSS2, KLK3, and the androgen receptor (AR). All four nodes are colored in blue, based on literature findings that show that the drug abiraterone has a downstream inhibitory effect on these molecules. Abiraterone acetate is a steroidal progesterone derivative with enhanced bioavailability used to treat refractory prostate cancer, as one of its mechanisms of action is the inhibition of CYP17A1, a steroidogenic enzyme is up-regulated during the transition of prostate cancer cell phenotype to hormone-resistant ^(56)^. Both nodes, CYP17A1 and AR, are related in the male sex hormone signaling. A recent study showed a correlation between increased androgen levels and susceptibility in male covid-19 patients and showed that inhibition of AR and its ligand is related to decreased angiotensin-converting enzyme 2 (ACE2) and TMPRSS2 protein levels, both enzymes related to SARS-CoV-2 spike-RBD internalization ^(57)^. TMPRSS2 is also a node connected to abiraterone acetate, as androgens and AR ^(58)^. A link between the AR, male sex hormones, and covid-19 severity has been proposed ^(59)^, and it was shown that abiraterone acetate exerts antiviral activity during the entry and post-entry events of SARS-CoV-2 replication cycle ^(55)^.

We also observed a modest antiviral activity of the aminoglycoside antibiotic neomycin against SARS-CoV-2. Antibiotics have been reported to inhibit eukaryotic translation ^(60, 61)^ directly, inhibit mitochondrial function ^(61–63)^, and induce changes in mammalian metabolic pathways. Aminoglycosides have been described to have other activities in addition to their antibacterial properties ^(64, 65)^. Prophylactic application of aminoglycosides to the nasal or vaginal mucosa of mice was shown to protect the animals from infection with DNA or RNA viruses ^(62)^. Aminoglycosides also induce the mucosal recruitment and expression of interferon-stimulated genes (ISGs) in dendritic cells in a Toll-like receptor 3 (TLR3)-dependent manner ^(65)^. Neomycin and kanamycin induce ISGs, while others such as streptomycin and amikacin did not. Interestingly, topical administration also increased mice survival after intranasal challenge with influenza and reduced viral shedding after intravaginal challenge with ZIKV virus ^(65)^. Our results showed that neomycin could decrease SARS-CoV-2 infection in Vero E6 cells.

Another interesting finding is that the extract of leaves of *Hedera helix* (common ivy) showed antiviral SARS-CoV-2 activity, decreasing the viral load on infected Vero E6 cells without showing cytotoxicity up to 50 μg/mL. This preparation has been clinically used to treat cough and showed beneficial effects for the treatment of inflammatory bronchial diseases ^(66)^. Moreover, the extract showed antiviral activity against the enterovirus EV71 C3 ^(67)^. In this study, the antiviral activity of hederasaponin B (a significant component of the extract) against EV71 C3 was verified and related to decreasing cytopathic effect (CPE) formation. In a mice model of influenza, coadministration of oseltamivir and *Hedera helix* extract resulted in increased protection of virus-infected mice, suggesting that *Hedera helix* extract enabled mice to overcome influenza virus infection when oseltamivir was given at sub-efficacious concentrations ^(68)^. This combination demonstrated benefits, including reduced lung inflammation and the potential to remove residual influenza viral particles. Network prediction analysis has shown that *Hedera helix* may inhibit FURIN activity through rutin and quercetin. It was previously shown that the insertion of a FURIN cleavage site in SARS-CoV, by genome mutation experiments, was responsible for its increased virulence and high capacity to enter into human lung cells in vitro. A recent study described that both SARS-CoV and SARS-CoV-2 uses the ACE2 receptor to facilitate viral entry into target cells. It was suggested that SARS-CoV-2 recognizes ACE2 more efficiently than SARS-CoV, increasing transmissibility ^(69)^. The ACE2 receptor is highly expressed in alveolar epithelial type II cells and is found at high levels in many extrapulmonary tissues. ACE2 is a component of the renin-angiotensin system, which regulates blood pressure. In the *Hedera helix* treated network, we observed that ACE2 is predicted to be inhibited. Our data showed that the extract of *Hedera helix* should be explored further as a source of antiviral compounds that could be used as adjuvant therapy against SARS-CoV-2 and other viral infections. Considering the strong connections between the flavonoids rutin and quercetin with several elements within the network associated with antivirus responses, it is relevant to highlight that not only *H. helix* but also other species displaying both flavonoids are used against influenza as folk medicine. Like *H. helix*, some species from the genus *Micania* (popularly known in Brazil as guaco and also sold in popular markets) displays quercetin in its composition ^(70)^ and are widely used in the country to treat colds, flu, and asthma, especially in poor rural and urban settings.

## Conclusion

A combination of phenotypic screening and network analysis provided new insights and starting points for the clinical evaluation and repurposing of drugs to inhibit SARS-CoV-2 infection and treat covid-19 patients at different disease stages. We have identified two drugs with potent and selective antiviral activity against SARS-CoV-2 *in vitro*, brequinar and abiraterone acetate, and two drugs with moderate activity, neomycin, and the extract of *Hedera helix*, that could be further explored as therapeutic adjuvants or as starting points for drug discovery for covid-19.

## Acknowledgments

The authors would like to thank Eurofarma Laboratórios for providing compounds and the researchers from the Molecular and Clinical Virology from ICB-USP for providing the SARS-CoV-2 isolate used in this study. DFSM received a scholarship from the Nacional Council for Scientific and Technological Development (CNPq). LMDR and KRG received postdoctoral fellowships from the Drugs for Neglected Diseases *initiative* through the Foundation for the Technological Development of Engineering from the University of Sao Paulo (Grant No. 2019/1913). This study was funded by the Sao Paulo Research Foundation (FAPESP - Grant No. 2020/04602-9).

---

**Figure S1.**
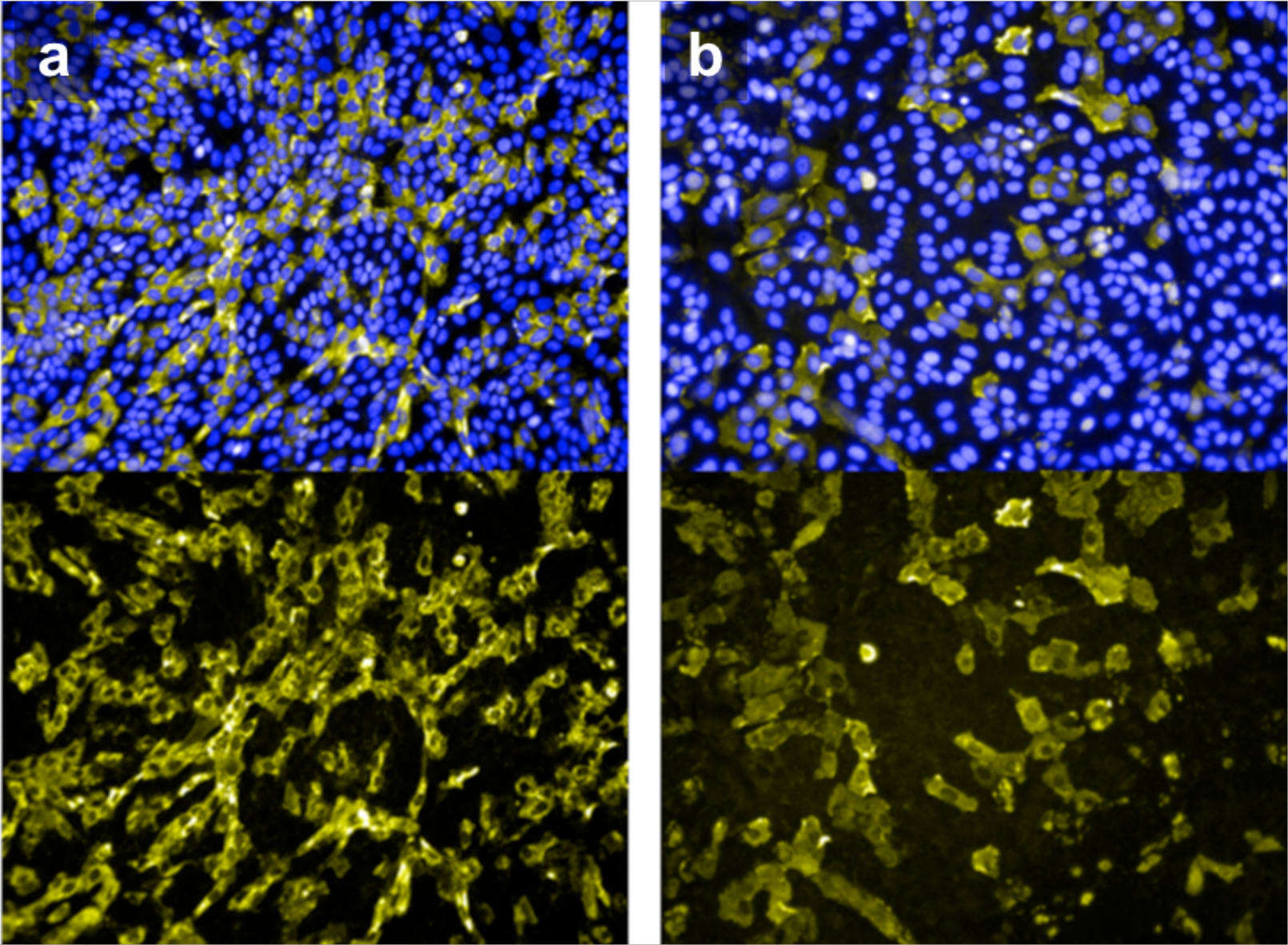
Comparison between the use for immunofluorescence experiments of: (*a*) A human sera isolated from SARS-CoV-2 convalescent patient and (b) a commercial polyclonal rabbit antibody against SARS-CoV-2 nucleoprotein.

## References

1. Yang, X., Yu, Y., Xu, J., Shu, H., Liu, H., Wu, Y., Wang, Y. et al. (2020). Clinical course and outcomes of critically ill patients with SARS-CoV-2 pneumonia in Wuhan, China: a single-centered, retrospective, observational study. The Lancet Respiratory Medicine doi: https://doi.org/10.1016/S2213-2600(20)30079-5

2. Riva, L., Yuan, S., Yin, X., Martin-Sancho, L., Matsunaga, N., Burgstaller, S., Chan, J. F. et al. (2020). A Large-scale Drug Repositioning Survey for SARS-CoV-2 Antivirals. bioRxiv. doi: https://doi.org/10.1101/2020.04.16.044016

3. World_Health_Organization. Accessed in 19/June/2020 https://covid19.who.int

4. Su, S., Wong, G., Shi, W., Liu, J., Lai, A. C., Zhou, J., Gao, G. F. et al. (2016). Epidemiology, genetic recombination, and pathogenesis of coronaviruses. Trends in microbiology, 24(6), 490–502. doi: https://doi.org/10.1016/j.tim.2016.03.003

5. De Wit, E., Van Doremalen, N., Falzarano, D., & Munster, V. J. (2016). SARS and MERS: recent insights into emerging coronaviruses. Nature Reviews Microbiology, 14(8), 523. doi: http://doi.org/10.1038/nrmicro.2016.81

6. Gordon, D. E., Jang, G. M., Bouhaddou, M., Xu, J., Obernier, K., White, K. M., Tummino, T. A. et al. (2020). A SARS-CoV-2 protein interaction map reveals targets for drug repurposing. Nature, 1–13. doi: https://doi.org/10.1038/s41586-020-2286-9

7. Anderson, R. M., Heesterbeek, H., Klinkenberg, D., & Hollingsworth, T. D. (2020). How will country-based mitigation measures influence the course of the COVID-19 epidemic? The Lancet, 395(10228), 931–934. doi: https://doi.org/10.1016/S0140-6736(20)30567-5

8. Gates, B. (2020). Responding to Covid-19—a once-in-a-century pandemic? New England Journal of Medicine, 382(18), 1677–1679. doi: https://doi.org/10.1056/NEJMp2003762

9. Horby, P., Lim, W. S., Emberson, J., Mafham, M., Bell, J., Linsell, L., Prudon, B. et al. (2020). Effect of Dexamethasone in Hospitalized Patients with COVID-19: Preliminary Report. medRxiv. doi: https://doi.org/10.1101/2020.06.22.20137273

10. Wang M, Cao R, Zhang L, Yang X, Liu J, Xu M, et al. (2020. Remdesivir and chloroquine effectively inhibit the recently emerged novel coronavirus (2019-nCoV) in vitro. Cell Res. 30: 269–271. doi: https://doi.org/10.1038/s41422-020-0282-0

11. Liu, J., Cao, R., Xu, M., Wang, X., Zhang, H., Hu, H., Wang, M. et al. (2020). Hydroxychloroquine, a less toxic derivative of chloroquine, is effective in inhibiting SARS-CoV-2 infection in vitro. Cell discovery, 6(1), 1–4. doi: https://doi.org/10.1038/s41421-020-0156-0

12. Caly, L., Druce, J. D., Catton, M. G., Jans, D. A., & Wagstaff, K. M. (2020). The FDA-approved drug ivermectin inhibits the replication of SARS-CoV-2 in vitro. Antiviral research, 104787. doi: https://doi.org/10.1016/j.antiviral.2020.104787

13. Touret, F., Gilles, M., Barral, K., Nougairède, A., Decroly, E., de Lamballerie, X., & Coutard, B. (2020). In vitro screening of a FDA approved chemical library reveals potential inhibitors of SARS-CoV-2 replication. BioRxiv. doi: https://doi.org/10.1101/2020.04.03.023846

14. Choy, K. T., Wong, A. Y. L., Kaewpreedee, P., Sia, S. F., Chen, D., Hui, K. P. Y., Peiris, M. et al. (2020). Remdesivir, lopinavir, emetine, and homoharringtonine inhibit SARS-CoV-2 replication in vitro. Antiviral research, 104786. doi: https://10.1016/j.antiviral.2020.104786

15. Shannon, A., Selisko, B., Le, N. T. T., Huchting, J., Touret, F., Piorkowski, G., Coutard, B. et al. (2020). Favipiravir strikes the SARS-CoV-2 at its Achilles heel, the RNA polymerase. bioRxiv doi: https://doi.org/10.1101%2F2020.05.15.09873116.

16. Wang, X., Cao, R., Zhang, H., Liu, J., Xu, M., Hu, H., Yang, X. et al. (2020). The antiinfluenza virus drug, arbidol is an efficient inhibitor of SARS-CoV-2 in vitro. Cell Discovery, 6(1), 1–5. https://doi.org/10.1038/s41421-020-0169-8

17. Zhou, Y., Hou, Y., Shen, J., Huang, Y., Martin, W., & Cheng, F. (2020). Network-based drug repurposing for novel coronavirus 2019-nCoV/SARS-CoV-2. Cell discovery, 6(1), 1–18. doi: https://doi.org/10.1038/s41421-020-0153-3

18. Bobrowski, T., Chen, L., Eastman, R. T., Itkin, Z., Shinn, P., Chen, C., Hall, M. et al. (2020). Discovery of Synergistic and Antagonistic Drug Combinations against SARS-CoV-2 In Vitro. bioRxiv. doi: https://doi.org/10.1101/2020.06.29.178889

19. Park JG, Ávila-Pérez G, Nogales A, Blanco-Lobo P, de la Torre JC, Martínez-Sobrido L. Identification and Characterization of Novel Compounds with Broad-Spectrum Antiviral Activity against Influenza A and B Viruses. J Virol. 2020;94(7):e02149–19. Published 2020 Mar 17. doi: https://doi.org/10.1128/JVI.02149-19

20. Qing, M., Zou, G., Wang, Q. Y., Xu, H. Y., Dong, H., Yuan, Z., & Shi, P. Y. (2010). Characterization of dengue virus resistance to brequinar in cell culture. Antimicrobial agents and chemotherapy, 54(9), 3686–3695. doi: https://doi.org/10.1128/AAC.00561-10

21. Xiong, R., Zhang, L., Li, S., Sun, Y., Ding, M., Wang, Y., Shan, J. et al. (2020). Novel and potent inhibitors targeting DHODH, a rate-limiting enzyme in de novo pyrimidine biosynthesis, are broad-spectrum antiviral against RNA viruses including newly emerged coronavirus SARS-CoV-2. BioRxiv. doi: https://doi.org/10.1101/2020.03.11.983056

22. Pilger, D. R., Moraes, C. B., Gil, L. H., & Freitas-Junior, L. H. (2017). Drug repurposing for yellow fever using high content screening. BioRxiv, 225656. doi: https://doi.org/10.1101/225656

23. Pascoalino, B. S., Courtemanche, G., Cordeiro, M. T., Gil, L. H., & Freitas-Junior, L. (2016). Zika antiviral chemotherapy: identification of drugs and promising starting points for drug discovery from an FDA-approved library. F1000Research, 5. doi: https://doi.org/10.12688/f1000research.9648.1

24. Park, J. G., Ávila-Pérez, G., Nogales, A., Blanco-Lobo, P., Juan, C., & Martínez-Sobrido, L. (2020). Identification and Characterization of Novel Compounds with Broad-Spectrum Antiviral Activity against Influenza A and B Viruses. Journal of Virology, 94(7). doi: https://doi.org/10.1128/JVI.02149-19

25. Ghosh, A., Nayak, R., & Shaila, M. S. (1996). Inhibition of replication of rinderpest virus by 5-fluorouracil. Antiviral research, 31(1-2), 35–44. doi: https://doi.org/10.1016/0166-3542(96)00943-6

26. Mingorance, L., Friesland, M., Coto-Llerena, M., Pérez-del-Pulgar, S., Boix, L., López-Oliva, J. M., Gastaminza, P. et al. (2014). Selective inhibition of hepatitis C virus infection by hydroxyzine and benztropine. Antimicrobial agents and chemotherapy, 58(6), 3451–3460. doi: https://doi.org/10.1128/aac.02619-14

27. Nawa, M. (1998). Effects of bafilomycin Aff1 on Japanese encephalitis virus in C6/36 mosquito cells. Archives of virology, 143(8), 1555–1568. doi: https://doi.org/10.1007/s007050050398

28. Yeganeh, B., Ghavami, S., Kroeker, A. L., Mahood, T. H., Stelmack, G. L., Klonisch, T., Halayko, A. J. et al. (2015). Suppression of influenza A virus replication in human lung epithelial cells by non-cytotoxic concentrations bafilomycin A1. American Journal of Physiology-Lung Cellular and Molecular Physiology, 308(3), L270–L286. doi: https://doi.org/10.1152/ajplung.00011.2014

29. Boulware, D. R., Pullen, M. F., Bangdiwala, A. S., Pastick, K. A., Lofgren, S. M., Okafor, E. C., Engen, N. W. et al. (2020). A randomized trial of hydroxychloroquine as postexposure prophylaxis for Covid-19. New England Journal of Medicine. doi: https://doi.org/10.1056/NEJMoa2016638

30. Chen, L., Zhang, Z. Y., Fu, J. G., Feng, Z. P., Zhang, S. Z., Han, Q. Y., Chen, X. L. et al. (2020). Efficacy and safety of chloroquine or hydroxychloroquine in moderate type of COVID-19: a prospective open-label randomized controlled study. medRxiv. doi: https://doi.org/10.1101/2020.06.19.20136093

31. Tang, W., Cao, Z., & Han, M. Hydroxychloroquine in patients mainly with mild to moderate COVID-19: an open-label, randomized, controlled trial. medRxiv 2020; published online May 7. doi: https://doi.org/10.1101/2020.04.10.20060558

32. Huang, M., Li, M., Xiao, F., Pang, P., Liang, J., Tang, T., Li, Y. et al. (2020). Preliminary evidence from a multicenter prospective observational study of the safety and efficacy of chloroquine for the treatment of COVID-19. National Science Review. doi: https://doi.org/10.1093/nsr/nwaa113

33. Geleris, J., Sun, Y., Platt, J., Zucker, J., Baldwin, M., Hripcsak, G., Sobieszczyk, M. E. et al. (2020). Observational study of hydroxychloroquine in hospitalized patients with Covid-19. New England Journal of Medicine. doi: https://doi.org/10.1056/NEJMoa2012410

34. Saleh, M., Gabriels, J., Chang, D., Kim, B. S., Mansoor, A., Mahmood, E., Beldner, S. et al. (2020). The effect of chloroquine, hydroxychloroquine and azithromycin on the corrected QT interval in patients with SARS-CoV-2 infection. Circulation: Arrhythmia and Electrophysiology. doi: https://doi.org/10.1161/circep.120.008662

35. La Frazia, S., Ciucci, A., Arnoldi, F., Coira, M., Gianferretti, P., Angelini, M., Santoro, M. G. et al. (2013). Thiazolides, a new class of antiviral agents effective against rotavirus infection, target viral morphogenesis, inhibiting viroplasm formation. Journal of virology, 87(20), 11096–11106. doi: https://doi.org/10.1128%2FJVI.01213-13

36. Meneses, M. D. S., Duarte, R. S., Migowski, E. R., & Ferreira, D. F. (2013). In vitro study on the effects of nitazoxanide on the replication of dengue virus and yellow fever virus. In Poster Presented at the 28th International Conference on Antiviral Research (ICAR). Abstract (Vol. 157, p. 101). doi: https://doi.org/10.1016/j.antiviral.2014.07.014

37. Shi, Z., Wei, J., Deng, X., Li, S., Qiu, Y., Shao, D., Ma, Z. et al. (2014). Nitazoxanide inhibits the replication of Japanese encephalitis virus in cultured cells and in a mouse model. Virology journal, 11(1), 10. doi: https://doi.org/10.1186/1743-422X-11-10

38. Rossignol, J. F. (2016). Nitazoxanide, a new drug candidate for the treatment of Middle East respiratory syndrome coronavirus. Journal of infection and public health, 9(3), 227–230. doi: https://doi.org/10.1016/j.jiph.2016.04.001

39. Jasenosky, L. D., Cadena, C., Mire, C. E., Borisevich, V., Haridas, V., Ranjbar, S., Cassell, G. H. et al. (2019). The FDA-approved oral drug nitazoxanide amplifies host antiviral responses and inhibits Ebola virus. iScience, 19, 1279–1290. doi: https://doi.org/10.1016/j.isci.2019.07.003

40. Caly, L., Druce, J. D., Catton, M. G., Jans, D. A., & Wagstaff, K. M. (2020). The FDA-approved drug ivermectin inhibits the replication of SARS-CoV-2 in vitro. Antiviral research, 104787. doi: https://doi.org/10.1016/j.antiviral.2020.104787

41. Patel, A., & Desai, S. (2020). Ivermectin in COVID-19 related critical illness. Available at SSRN 3570270.

42. Mastrangelo, E., Pezzullo, M., De Burghgraeve, T., Kaptein, S., Pastorino, B., Dallmeier, K., Bolognesi, M. et al. (2012). Ivermectin is a potent inhibitor of flavivirus replication specifically targeting NS3 helicase activity: new prospects for an old drug. Journal of Antimicrobial Chemotherapy, 67(8), 1884–1894. doi: https://doi.org/10.1093%2Fjac%2Fdks147

43. Varghese, F. S., Kaukinen, P., Gläsker, S., Bespalov, M., Hanski, L., Wennerberg, K., Ahola, T. et al. (2016). Discovery of berberine, abamectin and ivermectin as antivirals against chikungunya and other alphaviruses. Antiviral research, 126, 117–124. doi: https://doi.org/10.1016/j.antiviral.2015.12.012

44. Nguyen, K. Y., Sakuna, K., Kinobe, R., & Owens, L. (2014). Ivermectin blocks the nuclear location signal of parvoviruses in crayfish, Cherax quadricarinatus. Aquaculture, 420, 288–294. doi: https://doi.org/10.1016/j.aquaculture.2013.11.022

45. Wagstaff, K. M., Sivakumaran, H., Heaton, S. M., Harrich, D., & Jans, D. A. (2012). Ivermectin is a specific inhibitor of importin α/β-mediated nuclear import able to inhibit replication of HIV-1 and dengue virus. Biochemical Journal, 443(3), 851–856. doi: https://doi.org/10.1042%2FBJ20120150

46. Pizzorno, G., Wiegand, R. A., Lentz, S. K., & Handschumacher, R. E. (1992). Brequinar potentiates 5-fluorouracil antitumor activity in a murine model colon 38 tumor by tissuespecific modulation of uridine nucleotide pools. Cancer research, 52(7), 1660–1665.

47. Dexter, D. L., Hesson, D. P., Ardecky, R. J., Rao, G. V., Tippett, D. L., Dusak, B. A., Forbes, M. et al. (1985). Activity of a novel 4-quinolinecarboxylic acid, NSC 368390 [6-fluoro-2-(2’-fluoro-1, 1’-biphenyl-4-yl)-3-methyl-4-quinolinecarboxylic acid sodium salt], against experimental tumors. Cancer research, 45(11 Part 1), 5563–5568

48. Wang, Q. Y., Bushell, S., Qing, M., Xu, H. Y., Bonavia, A., Nunes, S., Dong, H. et al. (2011). Inhibition of dengue virus through suppression of host pyrimidine biosynthesis. Journal of virology, 85(13), 6548–6556. doi: https://doi.org/10.1128/JVI.02510-10

49. Hoffmann, H. H., Kunz, A., Simon, V. A., Palese, P., & Shaw, M. L. (2011). Broad-spectrum antiviral that interferes with de novo pyrimidine biosynthesis. Proceedings of the National Academy of Sciences, 108(14), 5777–5782. doi: https://doi.org/10.1073/pnas.1101143108

50. Ortiz-Riaño, E., Ngo, N., Devito, S., Eggink, D., Munger, J., Shaw, M. L., Martínez-Sobrido, L. et al. (2014). Inhibition of arenavirus by A3, a pyrimidine biosynthesis inhibitor. Journal of virology, 88(2), 878–889. doi: https://doi.org/10.1128/JVI.02275-13

51. Wang, Y., Wang, W., Xu, L., Zhou, X., Shokrollahi, E., Felczak, K., Metselaar, H. J. et al. (2016). Cross talk between nucleotide synthesis pathways with cellular immunity in constraining hepatitis E virus replication. Antimicrobial agents and chemotherapy, 60(5), 2834–2848. doi: https://doi.org/10.1128%2FAAC.02700-15

52. Xiong, R., Zhang, L., Li, S., Sun, Y., Ding, M., Wang, Y., Shan, J. et al. (2020). Novel and potent inhibitors targeting DHODH, a rate-limiting enzyme in de novo pyrimidine biosynthesis, are broad-spectrum antiviral against RNA viruses including newly emerged coronavirus SARS-CoV-2. BioRxiv. doi: https://doi.org/10.1101/2020.03.11.983056

53. Luthra, P., Naidoo, J., Pietzsch, C. A., De, S., Khadka, S., Anantpadma, M., Ready, J. M. et al. (2018). Inhibiting pyrimidine biosynthesis impairs Ebola virus replication through depletion of nucleoside pools and activation of innate immune responses. Antiviral research, 158, 288–302. doi: https://doi.org/10.1016/j.antiviral.2018.08.012

54. Hadjadj, J., Yatim, N., Barnabei, L., Corneau, A., Boussier, J., Pere, H., Carlier, N. et al. (2020). Impaired type I interferon activity and exacerbated inflammatory responses in severe Covid-19 patients. MedRxiv. doi: https://doi.org/10.1101/2020.04.19.20068015

55. Yuan, S., Chan, J. F., Chik, K. K., Chan, C. C., Tsang, J. O., Liang, R., Cai, J. P. et al. (2020). Discovery of the FDA-approved drugs bexarotene, cetilistat, diiodohydroxyquinoline, and abiraterone as potential COVID-19 treatments with a robust two-tier screening system. Pharmacological Research, 104960. doi: https://doi.org/10.1016/j.phrs.2020.104960

56. Giatromanolaki, A., Fasoulaki, V., Kalamida, D., Mitrakas, A., Kakouratos, C., Lialiaris, T., & Koukourakis, M. I. (2019). CYP17A1 and Androgen-Receptor Expression in Prostate Carcinoma Tissues and Cancer Cell Lines. Current Urology, 13(3), 157–165. doi: https://doi.org/10.1159/000499276

57. Stopsack, K. H., Mucci, L. A., Antonarakis, E. S., Nelson, P. S., & Kantoff, P. W. (2020). TMPRSS2 and COVID-19: Serendipity or Opportunity for Intervention?. Cancer discovery, 10(6), 779–782. doi: https://doi.org/10.1158/2159-8290.CD-20-0451

58. Lin, B., Ferguson, C., White, J. T., Wang, S., Vessella, R., True, L. D., Nelson, P. S. et al. (1999). Prostate-localized and androgen-regulated expression of the membrane-bound serine protease TMPRSS2. Cancer research, 59(17), 4180–4184.

59. Goren, A., McCoy, J., Wambier, C. G., Vano-Galvan, S., Shapiro, J., Dhurat, R., Lotti, T. et al. (2020). What does androgenetic alopecia have to do with COVID-19? An insight into a potential new therapy. Dermatologic therapy. doi: https://doi.org/10.1111%2Fdth.13365

60. Badal, S., Her, Y. F., & Maher, L. J. (2015). Nonantibiotic effects of fluoroquinolones in mammalian cells. Journal of Biological Chemistry, 290(36), 22287–22297. doi: https://dx.doi.org/10.1074%2Fjbc.M115.671222

61. Gopinath, S., Kim, M. V., Rakib, T., Wong, P. W., van Zandt, M., Barry, N. A., Iwasaki, A. et al. (2018). Topical application of aminoglycoside antibiotics enhances host resistance to viral infections in a microbiota-independent manner. Nature microbiology, 3(5), 611–621. doi: https://doi.org/10.1038/s41564-018-0138-2

62. Kalghatgi, S., Spina, C. S., Costello, J. C., Liesa, M., Morones-Ramirez, J. R., Slomovic, S., Collins, J. J. et al. (2013). Bactericidal antibiotics induce mitochondrial dysfunction and oxidative damage in mammalian cells. Science translational medicine, 5(192), 192ra85–192ra85. doi: https://doi.org/10.1126/scitranslmed.3006055

63. Moullan, N., Mouchiroud, L., Wang, X., Ryu, D., Williams, E. G., Mottis, A., Houtkooper, R. H. et al. (2015). Tetracyclines disturb mitochondrial function across eukaryotic models: a call for caution in biomedical research. Cell reports, 10(10), 1681–1691. doi: https://doi.org/10.1016%2Fj.celrep.2015.02.034

64. Cohen, J. I. (2018). New activities for old antibiotics. Nature microbiology, 3(5), 531–532. doi: https://doi.org/10.1038/s41564-018-0152-4

65. Krause, K. M., Serio, A. W., Kane, T. R., & Connolly, L. E. (2016). Aminoglycosides: an overview. Cold Spring Harbor perspectives in medicine, 6(6), a027029. doi: https://doi.org/10.1101/cshperspect.a027029

66. Hocaoglu, A. B., Karaman, O., Erge, D. O., Erbil, G., Yilmaz, O., Kivcak, B., Uzuner, N. et al. (2012). Effect of Hedera helix on lung histopathology in chronic asthma. Iranian Journal of Allergy, Asthma and Immunology, 316–323.

67. Song, J., Yeo, S. G., Hong, E. H., Lee, B. R., Kim, J. W., Kim, J., Park, J. H. et al. (2014). Antiviral activity of hederasaponin B from hedera helix against enterovirus 71 subgenotypes C3 and C4a. Biomolecules & Therapeutics, 22(1), 41. doi: https://doi.org/10.4062%2Fbiomolther.2013.108

68. Hong, E. H., Song, J. H., Shim, A., Lee, B. R., Kwon, B. E., Song, H. H., Seo, S. U. et al. (2015). Co-administration of Hedera helix L. extract enabled mice to overcome insufficient protection against influenza A/PR/8 virus infection under suboptimal treatment with oseltamivir. PloS one, 10(6), e0131089. doi: https://doi.org/10.1371/journal.pone.0131089

69. Yan, R., Zhang, Y., Li, Y., Xia, L., Guo, Y., & Zhou, Q. (2020). Structural basis for the recognition of SARS-CoV-2 by full-length human ACE2. Science, 367(6485), 1444–1448. doi: https://doi.org/10.1126/science.abb2762

70. Napimoga, M. H., & Yatsuda, R. (2010). Scientific evidence for Mikania laevigata and Mikania glomerata as a pharmacological tool. Journal of Pharmacy and Pharmacology, 62(7), 809–820. doi: https://doi.org/10.1211/jpp.62.07.0001

