## Supplemental Table 1 for "Discovery of clinically approved drugs capable of inhibiting SARS-CoV-2 *in vitro* infection using a phenotypic screening strategy and network-analysis to predict their potential to treat covid-19"

| Compoud Name | EC50 (M) | CC50 (M) |
|---|---|---|
| Abiraterone Acetate micronized | 7,10E-06 | > 50E-06 |
| Cefdinir | > 50E-06 | > 50E-06 |
| Cyanocobalamin | > 50E-06 | > 50E-06 |
| Ciprofloxacin | > 50E-06 | 6,51E-04 |
| Fexofenadine hydrochloride | 9,53E-05 | > 50E-06 |
| Ondasetron Hydrochloride | > 50E-06 | > 50E-06 |
| Erlotinib (Hydrochloride) | > 50E-06 | 2,91E-05 |
| Etodolac | > 50E-06 | > 50E-06 |
| Ibuprofen Dc90 | > 50E-06 | > 50E-06 |
| Levocetirizine Dihydrochloride | > 50E-06 | > 50E-06 |
| Minoxidil | > 50E-06 | > 50E-06 |
| Montelukast sodium | ND | ND |
| Nimesulide | > 50E-06 | > 50E-06 |
| Nimesulide Betacyclodextrin | > 50E-06 | > 50E-06 |
| Paracetamol Dc90 | > 50E-06 | > 50E-06 |
| Sugammadex | > 50E-06 | > 50E-06 |
| Tadalafil | > 50E-06 | > 50E-06 |
| Pyridoxine hydrochloride | > 50E-06 | > 50E-06 |
| Levetiracetam | > 50E-06 | > 50E-06 |
| Rivaroxaban | > 50E-06 | > 50E-06 |
| Letrozole | > 50E-06 | > 50E-06 |
| Tolnaftate | 2,26E-05 | 1,76E-05 |
| Ketoconazole | 1,20E-05 | 2,00E-05 |
| Fosfomycin trometamol | > 50E-06 | 8,93E-08 |
| Ketoprofen | > 50E-06 | > 50E-06 |
| Betamethasone | 4,90E-05 | 5,16E-04 |
| Zolpidem Hemitartrate | > 50E-06 | > 50E-06 |
| Micronized Ebastine | 5,74E-05 | 3,09E-06 |
| Olmesartan Medoxomil | > 50E-06 | > 50E-06 |
| Spironolactone | > 50E-06 | 1,51E-04 |
| Aceclofenac | 5,04E-04 | 1,38E-04 |
| Chondroitin Sulfate | > 50E-06 | > 50E-06 |
| Hydrochlorothiazide | > 50E-06 | > 50E-06 |
| Prednisolone Base | > 50E-06 | > 50E-06 |
| Atenolol | > 50E-06 | > 50E-06 |
| Potassium V. Penicilin | > 50E-06 | > 50E-06 |
| Glycosamine Sulfate | > 50E-06 | > 50E-06 |
| Doxazosin mesylate | 2,24E-05 | 3,50E-05 |
| Calcium Atorvastatin | 2,19E-06 | 3,71E-06 |
| Indapamide | > 50E-06 | > 50E-06 |
| Chlortalidone | > 50E-06 | 6,86E-04 |
| Gentamicin Sulfate | > 50E-06 | > 50E-06 |
| Silybum Marianum Dry Extract | > 50E-06 | > 50E-06 |
| Amiloride Hydrochloride | > 50E-06 | > 50E-06 |
| Benzathine Penicillin G | > 50E-06 | > 50E-06 |
| Hedera helix (g/mL) | 5,10E-04 | > 50E-06 |
| Neomycin sulfate | 1,29E-05 | > 50E-06 |
| Diosmin | > 50E-06 | 1,93E-07 |
| Triancinolone | > 50E-06 | > 50E-06 |
| Amoxicilin | > 50E-06 | 8,59E-08 |
| Azithromycin | > 50E-06 | > 50E-06 |
| Acetylsalicylic acid | > 50E-06 | > 50E-06 |
| Axetylcefuroxime | > 50E-06 | > 50E-06 |
| Mometasone Furoate | 7,34E-06 | 1,95E-05 |
| Budesonide | > 50E-06 | > 50E-06 |
| Tamoxifen | 7,81E-06 | 6,88E-05 |
