## Supplemental Table 2 for "Discovery of clinically approved drugs capable of inhibiting SARS-CoV-2 *in vitro* infection using a phenotypic screening strategy and network-analysis to predict their potential to treat covid-19"

| Symbol | Synonym(s) | Entrez Gene Name | Location | Family | Entrez Gene ID for Human | Entrez Gene ID for Mouse | Entrez Gene ID for Rat |
|---|---|---|---|---|---|---|---|
| ABCA1 | ABC-1, ATP-binding cassette, sub-family A (ABC1), member 1, ATP binding cassette subfamily A member 1, CERP, HDLCQTL13, HDLDT1, HPALP1, TGD | ATP binding cassette subfamily A member 1 | Plasma Membrane | transporter | 19 | 11303 | 313210 |
| Abcb1b | Abcb1, ATP-binding cassette, sub-family B (MDR/TAP), member 1B, mdr, Mdr1, Mdr1b, Pgy-1 | ATP-binding cassette, sub-family B (MDR/TAP), member 1B | Plasma Membrane | transporter |  | 18669 | 24646 |
| ABCG1 | ABC8, ATP binding cassette subfamily G member 1, AW413978, White, WHITE1 | ATP binding cassette subfamily G member 1 | Plasma Membrane | transporter | 9619 | 11307 | 85264 |
| ACE2 | 2010305L05Rik, ACEH, angiotensin I converting enzyme 2, angiotensin I converting enzyme (peptidyl-dipeptidase A) 2 | angiotensin I converting enzyme 2 | Plasma Membrane | peptidase | 59272 | 70008 | 302668 |
| AGT | AGTN, AI265500, ANG, angiotensinogen, angiotensinogen (serpin peptidase inhibitor, clade A, member 8), ANHU, ANRT, Aogen, hFLT1, PAT, SERPINA8 | angiotensinogen | Extracellular Space | growth factor | 183 | 11606 | 24179 |
| AGTR1 | 1810074K20Rik, AG2S, Agt1ar, AGTR1A, AGTR1B, AI551199, Angiotensin II receptor type 1, angiotensin II receptor, type 1a, Angtr-1a, AT1, AT1A, At1ar, AT1B, AT1BR, AT1R, AT1 receptor, AT2R1, AT2R1A, ATR1a, HAT1R | angiotensin II receptor type 1 | Plasma Membrane | G-protein coupled receptor | 185 | 11607 | 24180 |
| AKR1B1 | ADR, Ahr-1, Akr1b3, Akr1b4, aldo-keto reductase family 1 member B, aldo-keto reductase family 1, member B3 (aldose reductase), Aldolase Reductase, Aldor1, Aldose Reductase, ALDR1, ALDRED, Alr, ALR2, ALR-P-I, Androgen-induced aldose reductase, AR, RATALDRED | aldo-keto reductase family 1 member B | Cytoplasm | enzyme | 231 | 11677 | 24192 |
| ALB | Alb-1, Albumin, Albumin 1, albumin isomer 1, albumin isomer 2, albumin isomer 3, Albza, BCL002, HSA, PRO0883, PRO0903, PRO1341, PRO2044, SERUM ALBUMIN, serum albumin chain A, Serum albumin precursor | albumin | Extracellular Space | transporter | 213 | 11657 | 24186 |
| Angiotensin II receptor type 1 | AGTR1, At1r |  | Plasma Membrane | group |  |  |  |
| ANGPT2 | AGPT2, ANG2, ANG II, angiopoietin 2, Angiotensin 2, Angiotensin II, Angrp | angiopoietin 2 | Extracellular Space | growth factor | 285 | 11601 | 89805 |
| APLN | 6030430G11Rik, APEL, Apelin, Preproapelin, XNPEP2 | apelin | Extracellular Space | other | 8862 | 30878 | 58812 |
| AR | AIS, Andr, androgen receptor, AW320017, DHTR, HUMARA, HYSP1, KD, NR3C4, SBMA, SMAX1, Testosterone receptor, TFM | androgen receptor | Nucleus | ligand-dependent nuclear receptor | 367 | 11835 | 24208 |
| ATG5 | 2010107M05Rik, 3110067M24Rik, APG5, APG5L, APG5-LIKE, APOPTOSIS SPECIFIC, ASP, Atg5l, autophagy related 5, AW319544, C88337, hAPG5, PADDY, POT1, SCAR25 | autophagy related 5 | Cytoplasm | other | 9474 | 11793 |  |
| BAX | Bcl2-associated X, BCL2 associated X, apoptosis regulator, BCL2-associated X protein, BCL2L4 | BCL2 associated X, apoptosis regulator | Cytoplasm | transporter | 581 | 12028 | 24887 |
| BCL2L1 | bBclxl, BCL2L, BCL2-like 1, BCLX, Bcl-X beta, Bclx gamma, BCL-XL/S, Bcl-X β, Bclx γ, PPP1R52 | BCL2 like 1 | Cytoplasm | other | 598 | 12048 | 24888 |

| | | | | | | | |
|---|---|---|---|---|---|---|---|
| CASP3 | A830040C14Rik, AC-3, Caspase-3, CASPASE-3 p20, CC3, CPP-32, CPP32B, CPP32-beta, CPP32-β, Ice-like cysteine protease, Lice, mldy, SCA-1, YAMA | caspase 3 | Cytoplasm | peptidase | 836 | 12367 | 25402 |
| CASP8 | ALPS2B, CAP4, Caspase-8, FLICE, MACH, MCH5, PROCASP8 | caspase 8 | Nucleus | peptidase | 841 | 12370 | 64044 |
| CASP9 | AI115399, APAF-3, AW493809, Casp9 v1, Caspase-9, ICE-LAP6, MCH6, PPP1R56 | caspase 9 | Cytoplasm | peptidase | 842 | 12371 | 58918 |
| CCN2 | AMPHIROGULIN, cellular communication network factor 2, CTGF, CTGF isoform 1, CTGRP, Fibroblast-inducible secreted, Fisp12, HCS24, IGFBP8, IGFBP-RP2, NOV2, Tissue growth factor | cellular communication network factor 2 | Extracellular Space | growth factor | 1490 | 14219 | 64032 |
| CCR5 | AM4-7, CC-CKR-5, CCL4 receptor, CCL5 receptor, C-C motif chemokine receptor 5, C-C motif chemokine receptor 5 (gene/pseudogene), CD195, CHEMOKINE CC5 receptor, chemokine (C-C motif) receptor 5, CKR-5, CMKBR5, IDDM22, LOC727797 | C-C motif chemokine receptor 5 (gene/pseudogene) | Plasma Membrane | G-protein coupled receptor | 1234 | 12774 | 117029 |
| CD200 | CD200 antigen, CD200 molecule, Cspmo2, MOX1, MOX2, MRC, MRCOX2, OX-2 | CD200 molecule | Plasma Membrane | other | 4345 | 17470 | 24560 |
| Cd64 | FC gamma R1, Fc-gamma receptor 1, FcgammaRI, FC γ R1, Fc-γ receptor 1, FcγRI | | Other | group | | | |
| CEACAM1 | AI267129, bb-1, BGP, BGP1, Bgp2, BGPA, Bgpd, BGPI, BGPR, carcinoembryonic antigen-related cell adhesion molecule 1, carcinoembryonic antigen-related cell adhesion molecule 2, Cc1, C-CAM, CCAM105, CD66a, Cea-1, Cea-7, CEACAM1-4L, Ceacam2, Ceacam2-L, CEA cell adhesion molecule 1, ecto-atpase, Ha4, HV2, mCEA1, Mhv-1, MHVR, MHVR1, mmCGM1, mmCGM1a, mmCGM2, Pp120 | CEA cell adhesion molecule 1 | Plasma Membrane | transporter | 634 | 26365\|26367 | 81613 |
| CSN2 | Beta casein, CASB, Casein, casein beta, Casein β, CSNB, β-Casein | casein beta | Extracellular Space | kinase | 1447 | 12991 | 29173 |
| CST3 | ARMD11, CYSC, cystatin C, gamma-TRACE, HEL-S-2, γ-TRACE | cystatin C | Extracellular Space | other | 1471 | 13010 | 25307 |
| CXCL10 | C7, chemokine (C-X-C motif) ligand 10, CRG-2, C-X-C motif chemokine ligand 10, gamma-IFN INDUCIBLE EARLY RESPONSE, gIP-10, IFI10, IFNG INDUCIBLE protein-10, INP10, Interferon-inducible protein-10, IP-10, mob-1, SCYB10, SMALL INDUCIBLE CYTOKINE SUBFAMILY B (Cys-X-Cys), member 10, γ-IFN INDUCIBLE EARLY RESPONSE | C-X-C motif chemokine ligand 10 | Extracellular Space | cytokine | 3627 | 15945 | 245920 |
| Cxcl11 | betaR1, b-R1, chemokine (C-X-C motif) ligand 11, Cxc11, H174, Ip9, I-tac, LOC630447, Scyb11, Scyb9b | chemokine (C-X-C motif) ligand 11 | Extracellular Space | cytokine | | 56066 | |
| CXCL3 | chemokine (C-X-C motif) ligand 2, Cinc-2, CINC-2a, CINC-2b, Cinc3, Cxcl2, C-X-C motif chemokine ligand 2, C-X-C motif chemokine ligand 3, Dcip1, Gm1960, GRO1, Gro2, GRO3, GROA, Gro-alpha, GROb, Gro beta, GROg, Gro-α, Gro β, Gro γ, Mgsa-b, MIP-2, MIP-2a, Mip2 alpha, MIP-2b, Mip2 α, Scyb, Scyb15, Scyb2, SCYB3 | C-X-C motif chemokine ligand 3 | Extracellular Space | cytokine | 2921 | 20310 | 114105 |
| CXCL8 | C-X-C motif chemokine ligand 8, GCP-1, IL8, LECT, LUCT, LYNAP, MDNCF, MONAP, Monocyte-derived neutrophil chemotactic factor, NAF, NAP-1 | C-X-C motif chemokine ligand 8 | Extracellular Space | cytokine | 3576 | | |

| | | | | | | | |
|---|---|---|---|---|---|---|---|
| CYP17A1 | 17 alpha-hydroxylase, 17 α-hydroxylase, C17,20 Lyase, CPT7, CYP17, CYP17A, CYPXVII, cytochrome P450 family 17 subfamily A member 1, cytochrome P450, family 17, subfamily a, polypeptide 1, p450 17alpha, p450 17alpha hydroxylase, P450C17, P450c17 alpha, P450c17 α, S17AH | cytochrome P450 family 17 subfamily A member 1 | Cytoplasm | enzyme | 1586 | 13074 | 25146 |
| CYP3A4 | CP33, CP34, CYP3A, Cyp3a11, Cyp3a2, CYP3A23, CYP3A3, CYPIIIA3, CYPIIIA4, CYTOCHROME P450 3A3, cytochrome P450 family 3 subfamily A member 4, cytochrome P450, family 3, subfamily a, polypeptide 2, HLP, NF-25, P450C3, P450PCN1, P-450ut-a | cytochrome P450 family 3 subfamily A member 4 | Cytoplasm | enzyme | 1576 | | 266682 |
| D-glucose | 110187-42-3, 13C-glucose, 2280-44-6, (3R,4S,5S,6R)-6-(hydroxymethyl)oxane-2,3,4,5-tetrol, 50-99-7, biliary glucose, C6H12O6, dextrose, D-glucose-13C6, glucose, sugar, (U-13C6)-D-glucose, [U-13C] glucose, U-C glucose, uniformly-labeled [U-13C]glucose | | Other | chemical - endogenous mammalian | PubChem ID 5793 | | |
| DHODH | 2810417D19Rik, AI834883, DHOdehase, DHOH, dihydroorotate dehydrogenase, dihydroorotate dehydrogenase (quinone), POADS, URA1 | dihydroorotate dehydrogenase (quinone) | Cytoplasm | enzyme | 1723 | 56749 | 65156 |
| ELAVL1 | 2410055N02Rik, DKFZP667b083, ELAV1, ELAV (embryonic lethal, abnormal vision)-like 1 (Hu antigen R), ELAV like RNA binding protein 1, Hua, Hu antigen R, HUR, MelG, RGD:731215, RNA binding protein HuR, W91709 | ELAV like RNA binding protein 1 | Cytoplasm | other | 1994 | 15568 | 363854 |
| FN1 | cFn, CIG, E330027I09, ED-B, FIBNEC, Fibronectin, FIBRONECTIN 1, Fibronectin3 M1, Fibronectin i, FINC, FN, FN1 isoform 1, FNZ, GFND, GFND2, LETS, MSF, SMDCF | fibronectin 1 | Extracellular Space | enzyme | 2335 | 14268 | 25661 |
| FOXM1 | AA408308, AW554517, BB238854, D1Mgi56, Fkh16, FKHL16, FOCM1, forkhead box M1, FOXM1B, HFH-11, HFH-11B, HNF-3, INS-1, MPHOSPH2, MPM2, MPP-2, PIG29, TRIDENT, WIN | forkhead box M1 | Nucleus | transcription regulator | 2305 | 14235 | 58921 |
| FURIN | 9130404I01RIK, BASIC-AMINO-ACID-SPECIFIC FURIN, FUR, FURIN FROM PACE, furin (paired basic amino acid cleaving enzyme), furin, paired basic amino acid cleaving enzyme, PACE, PCSK3, SPC1 | furin, paired basic amino acid cleaving enzyme | Cytoplasm | peptidase | 5045 | 18550 | 54281 |
| GATA4 | ASD2, GATA binding protein 4, TACHD, TOF, VSD1 | GATA binding protein 4 | Nucleus | transcription regulator | 2626 | 14463 | 54254 |
| GHRL | 2210006E23Rik, des-octanoyl ghrelin, Ghr, GHRELIN, ghrelin and obestatin prepropeptide, m46, MTLRP, MTLRPAP, OBESTATIN | ghrelin and obestatin prepropeptide | Extracellular Space | growth factor | 51738 | 58991 | 59301 |
| HMGB1 | Ac2-008, high mobility group box 1, HMG-1, HMG3, p30, SBP-1 | high mobility group box 1 | Nucleus | transcription regulator | 3146 | | |
| HRAS | C-BAS/HAS, C-HA-RAS, C-HA-RAS1, C-H-RAS, c-rasHa, CTLO, HAMSV, HA-RAS, Harvey-ras, Harvey rat sarcoma virus oncogene, HRAS1, H-RASIDX, HRas proto-oncogene, GTPase, Kras2, RAS, RASH1, RAS HA, V-H-RAS | HRas proto-oncogene, GTPase | Plasma Membrane | enzyme | 3265 | 15461 | 293621 |

| | | | | | | | |
|---|---|---|---|---|---|---|---|
| IFITM1 | 1110036C17Rik, 9-27, CD225, DSPA2a, HUM927A, IFI17, IFI27SEP, IFIM1, IFM1 9-27, IFN9-27, interferon induced transmembrane protein 1, LEU13, Mil-2 | interferon induced transmembrane protein 1 | Plasma Membrane | transmembrane receptor | 8519 | | |
| IFITM2 | 1-8D, DSPA2c, fragilis3, Ifi 16, IFI1-8U, IFITM3L, IFITML, IFN1-8D, interferon induced transmembrane protein 2, Interferon induced transmembrane protein 3-like, mil-3 | interferon induced transmembrane protein 2 | Cytoplasm | other | 10581 | 80876 | 114709 |
| IFNAR1 | alpha CHAIN of type I IFNR, AVP, Ifar, IFN-alpha-beta-R, IFNalpha/betaR, Ifn-alpha/beta-receptor, IFN alpha/beta receptor 1, IFN-alpha-REC, IFNAR, IFNBR, IFN receptor CHAIN 1, IFN receptor type 1, IFN type 1 receptor, IFN-α-REC, IFN α/βR, IFN-α-β-R, Ifn-α/β-receptor, IFN α/β receptor 1, IFRC, Infar, interferon (alpha and beta) receptor 1, interferon alpha and beta receptor subunit 1, Interferon Receptor, interferon (α and β) receptor 1, interferon α and β receptor subunit 1, LOC284829, type 1 interferon receptor, Type I IFNR, Type I infr, α CHAIN of type I IFNR, β r1 | interferon alpha and beta receptor subunit 1 | Plasma Membrane | transmembrane receptor | 3454 | 15975 | 288264 |
| IFNB1 | IFB, IFF, IFNB, IFNB1A, IFN-beta, IFN-beta1, Ifn beta1b, IFN-β, IFN-β1, interferon beta 1, interferon beta 1, fibroblast, interferon β 1, interferon β 1, fibroblast, neoferon, recombinant human interferon β 1a, recombinantinterferon beta 1a, recombinantinterferon β 1a | interferon beta 1 | Extracellular Space | cytokine | 3456 | 15977 | 24481 |
| IFNG | IFG, IFI, IFNG2, IFN gamma, IFN type II, IFN-γ, INF-γ, interferon gamma, Interferon γ, type II INTERFERON, γ-ifn, γ interferon | interferon gamma | Extracellular Space | cytokine | 3458 | 15978 | 25712 |
| IFNLR1 | CRF2-12, CRF2/12, IFN LAMBDA R1, IFNLR, IFN λ R1, Il28r, IL-28R1, IL28RA, interferon lambda receptor 1, interferon, lambda receptor 1, interferon λ receptor 1, interferon, λ receptor 1, LICR2, RGD1562689 | interferon lambda receptor 1 | Plasma Membrane | transmembrane receptor | 163702 | 242700 | 362628 |
| IL1 | | | Extracellular Space | group | | | |
| IL10 | CSIF, GVHDS, IL10A, IL10X, interleukin 10, TGIF | interleukin 10 | Extracellular Space | cytokine | 3586 | 16153 | 25325 |
| IL1A | IL1, IL1-ALPHA, IL-1F1, IL-1 α, interleukin 1 alpha, interleukin-1 α, Interleukin-A | interleukin 1 alpha | Extracellular Space | cytokine | 3552 | 16175 | 24493 |
| IL1B | IL-1, IL1-BETA, IL-1F2, IL-1β, interleukin 1 beta, Interleukin 1 β, OAF, Osteoclast-Activating Factor, Pro-IL-1beta, Pro-IL-1β | interleukin 1 beta | Extracellular Space | cytokine | 3553 | 16176 | 24494 |
| IL6 | BSF-2, CDF, FDGI, HGF, HSF, IFNB2, IFN-beta-2, IFN beta 2A, IFN-β-2, IFN β 2A, ILg6, interleukin-6 | interleukin 6 | Extracellular Space | cytokine | 3569 | 16193 | 24498 |
| IL9 | HP40, interleukin 9, P40 | interleukin 9 | Extracellular Space | cytokine | 3578 | 16198 | 116558 |
| Ins1 | Ins2-rs1, insulin 1, Insulin I, Preproinsulin, proinsulin | insulin I | Extracellular Space | other | | 16333 | 24505 |
| Insulin | Ins, Ins1/2 | | Extracellular Space | group | | | |
| ITGAM | C3R, CD11B, CD11B/CD18, Complement Receptor 5, COMPLEMENT receptor type 3, CR3, CR3A, F730045J24Rik, integrin alpha M, integrin subunit alpha M, integrin subunit α M, Integrin α m, ITGAN, Ly-40, MAC-1, MAC1A, MO1A, SLEB6 | integrin subunit alpha M | Plasma Membrane | transmembrane receptor | 3684 | 16409 | 25021 |

| | | | | | | | |
|---|---|---|---|---|---|---|---|
| KLK3 | 0610007D04Rik, alpha-NGF, APS, Egfbp-1, Egfbp2, Egfbp-3, EGF-BP A, EGF-BP C, Epidermal Growth Factor-Binding Protein Type A, gamma-NGF, Gk27, hK3, HNP, KAL, KAL-B, kallikrein, kallikrein 1, kallikrein 1-related pepidase b4, kallikrein 1-related peptidase b1, kallikrein 1-related peptidase b11, kallikrein 1-related peptidase b16, kallikrein 1-related peptidase b21, kallikrein 1-related peptidase b22, kallikrein 1-related peptidase b24, kallikrein 1-related peptidase b27, kallikrein 1-related peptidase b3, kallikrein 1-related peptidase b5, kallikrein 1-related peptidase b8, kallikrein 1-related peptidase b9, kallikrein 1-related petidase b26, kallikrein related peptidase 3, Klk1, Klk11, Klk16, Klk1b1, Klk1b11, Klk1b16, Klk1b21, Klk1b22, Klk1b24, Klk1b26, Klk1b27, Klk1b3, Klk1b4, Klk1b5, Klk1b6, Klk1b8, Klk1b9, Klk21, KLK21L, Klk22, Klk24, Klk26, Klk27, KLK2A1, Klk5, Klk6, Klk8, Klk9, mGK-1, mGK-11, mGk-16, mGk-21, mGk-22, mGk-24, mGK-26, mGK-27, mGk-3, mGk-4, mGK-5, mGk-6, mGK-8, mGk-9, mK1, Nerve growth factor gamma, Nerve growth factor γ, Ngfa, Ngfg, Nrpn, PRECE-2, Prostate-Specific Antigen, Proteinase a, Proteinase d, Proteinase f, PSA, TADG14, Tissue Kallikrein, α-NGF, γ-NGF, γ-seminoprotein | kallikrein related peptidase 3 | Extracellular Space | peptidase | 354 | 13646\|13648\|16612\|16613\|16615\|16616\|16617\|16618\|16619\|16622\|16623\|16624\|18048\|18050 | |
| LCK | Hck-3, IMD22, Lck1, LCK proto-oncogene, Src family tyrosine kinase, Lcktkr, LSK, Lskt, lymphocyte protein tyrosine kinase, p56Lck, pp58lck, YT16 | LCK proto-oncogene, Src family tyrosine kinase | Cytoplasm | kinase | 3932 | 16818 | 313050 |
| MAPK1 | 9030612K14Rik, AA407128, AU018647, C78273, ERK, ERK-2, ERK42, ERT1, Mapk1,2, MAPK2, Mapk p42, mitogen-activated protein kinase 1, MITOGEN ACTIVATED protein KINASE 2, p38, p40, p40 HERAK, p41, p41mapk, P42, p42 Erk, P42MAPK, PRKM1, PRKM2 | mitogen-activated protein kinase 1 | Cytoplasm | kinase | 5594 | 26413 | 116590 |
| MAPK3 | ERK-1, ERT2, Esrk1, HS44KDAP, HUMKER1A, MAPK1, Mapkapk3, Mapk p44, mitogen-activated protein kinase 3, MNK1, MTAP2K, p44, p44 Erk, P44ERK1, P44MAPK, PRKM3 | mitogen-activated protein kinase 3 | Cytoplasm | kinase | 5595 | 26417 | 50689 |
| Mmp | Matrix Metalloproteinase, MATRIX METALLOPROTEINASES, mmpi, mmp (Matrix Metalloproteinase), Mt-mmp | | Nucleus | group | | | |
| MMP2 | CLG4, CLG4A, GelA, GELATINASE, Gelatinase A, matrix metallopeptidase 2, METALLOPROTEINASE 2, MMP-II, MONA, TBE-1 | matrix metallopeptidase 2 | Extracellular Space | peptidase | 4313 | 17390 | 81686 |
| MMP9 | AW743869, B/MMP9, CLG4B, COLLAGENASE type IV, Gelatinase B, GELB, GI 92-kda, MANDP2, matrix metallopeptidase 9, METALLOPROTEINASE 9, pro-MMP-9 | matrix metallopeptidase 9 | Extracellular Space | peptidase | 4318 | 17395 | 81687 |
| NCF1 | LOC652699, NCF1A, NCF-47K, neutrophil cytosolic factor 1, NOXO2, P47 NADPH subunit, p47-phox, SH3PXD1A | neutrophil cytosolic factor 1 | Cytoplasm | enzyme | 653361 | 17969 | 114553 |

| | | | | | | | |
|---|---|---|---|---|---|---|---|
| NFE2L2 | BM974200, HEBP1, IMDDHH, NRF2, nuclear factor, erythroid 2-like 2, nuclear factor, erythroid derived 2, like 2 | nuclear factor, erythroid 2 like 2 | Nucleus | transcription regulator | 4780 | 18024 | 83619 |
| NOS3 | 2310065A03Rik, ECNOS, eNOS, nitric oxide synthase 3, nitric oxide synthase 3, endothelial cell, nNOS | nitric oxide synthase 3 | Cytoplasm | enzyme | 4846 | 18127 | 24600 |
| NQO1 | AV001255, DHQU, DIA4, DTD, DT-diaphorase, NAD DT-diaphorase, Nadph dehydrogenase, NAD(P)H dehydrogenase, quinone 1, Nadph diaphorase, NAD(P)H quinone dehydrogenase 1, NAD(P)H:quinone oxidoreductase, Nadph Quinone Oxidoreductase-1, NMO1, NMOR, NMOR1, NMORI, Nqo, Ox-1, Qr, QR1, Quinone reductase | NAD(P)H quinone dehydrogenase 1 | Cytoplasm | enzyme | 1728 | 18104 | 24314 |
| NTS | 5033428E16RIK, Neuromedin N, neurotensin, NmN, NMN-125, NN, NT, NT/N, NTS1 | neurotensin | Extracellular Space | other | 4922 | 67405 | 299757 |
| P glycoprotein | MDR, P-gp | | Plasma Membrane | group | | | |
| P2RX4 | AI504491, AW555605, D5Ertd444e, P2X4, P2X4R, purinergic receptor P2X 4, purinergic receptor P2X, ligand-gated ion channel 4 | purinergic receptor P2X 4 | Plasma Membrane | ion channel | 5025 | 18438 | 29659 |
| P2RX7 | AI467586, P2X7, P2X(7), P2X7R, P2Z, Purinergic receptor, purinergic receptor P2X 7, purinergic receptor P2X, ligand-gated ion channel, 7 | purinergic receptor P2X 7 | Plasma Membrane | ion channel | 5027 | 18439 | |
| PCSK1 | BDP, BMIQ12, DCE, Dyn-converting enzyme, NEC1, PC1, PC3, Phpp-1, proprotein convertase subtilisin/kexin type 1, SPC3 | proprotein convertase subtilisin/kexin type 1 | Cytoplasm | peptidase | 5122 | 18548 | |
| PCSK6 | b2b2830Clo, C86343, PACE4, PAIRED BASIC AMINO ACID CLEAVING ENZYME 4, proprotein convertase subtilisin/kexin type 6, SPC4 | proprotein convertase subtilisin/kexin type 6 | Extracellular Space | peptidase | 5046 | 18553 | 25507 |
| phosphatidylinositol | phosphatidylinositide, phosphoinositol, phosphoionositides, PI | | Other | chemical - endogenous mammalian | | | |
| POR | 4933424M13Rik, CPR, Cyp, Cyp450r, CYPOR, Cyp reductase, Cytochrome P450 Oxidoreductase, cytochrome P450 reductase, NADH OXYDOREDUCTASE, Nadph Cytochrome C Reductase, NADPH CYT P450 REDUCTASE, NADPH DEPENDENT CYTOCHROME P450 REDUCTASE, NADPH Reductase, P450 (cytochrome) oxidoreductase, P450 Oxidoreductase, P450R, P450 Reductase | cytochrome p450 oxidoreductase | Cytoplasm | enzyme | 5447 | 18984 | 29441 |
| PRL | AV290867, Dlt, GHA1, Prl1a1, PRLB, PRLSD1, Prol, Prolactin, RATPRLSD1, RNPROL | prolactin | Extracellular Space | cytokine | 5617 | 19109 | 24683 |
| PTGS2 | Cox, COX-2, CYCLO-OXYGENASE 2, GRIPGHS, hCox-2, INDUCIBLE CYCLOOXYGENASE, PES-2, PGG/HS, PGHS-2, PGH synthase 2, Pgi2 synthase, Pgs2, Pgsi, PHS-2, PHS II, Prostaglandin endoperoxide synthase 2, TIS10 | prostaglandin-endoperoxide synthase 2 | Cytoplasm | enzyme | 5743 | 19225 | 29527 |
| Rbp | Retinal binding, Retinoic Acid Binding, RETINOL binding | | Cytoplasm | group | | | |
| reactive oxygen species | oxygen and reactive oxygen species, reactive oxygen metabolites, ROI, ROS | | Other | chemical toxicant | | | |
| RETN | ADSF, FIZZ3, HXCP1, RESISTIN, RETN1, RSTN, XCP1, Xcp4 | resistin | Extracellular Space | other | 56729 | 57264 | 246250 |

| | | | | | | | |
|---|---|---|---|---|---|---|---|
| SERPINE1 | beta MIGRATING PLAS ACTIVATOR, beta-MIGRATING PLASMINOGEN ACTIVATOR INHIBITOR I, PAI, PAI-1, PAI1A, Pai1aa, Planh, PLANH1, Plasminogen activator inhibitor 1, RATPAI1A, serine (or cysteine) peptidase inhibitor, clade E, member 1, SERPINE, serpin family E member 1, β MIGRATING PLAS ACTIVATOR, β-MIGRATING PLASMINOGEN ACTIVATOR INHIBITOR I | serpin family E member 1 | Extracellular Space | other | 5054 | 18787 | 24617 |
| TIMP1 | CLGI, EPA, EPO, HCI, Metalloproteinase inhibitor, TIMP, TIMP metallopeptidase inhibitor 1, Tissue Inhibitor of Metalloproteinase 1, TPA-S1 | TIMP metallopeptidase inhibitor 1 | Extracellular Space | cytokine | 7076 | 21857 | 116510 |
| TLR2 | CD282, Ly105, TIL4, toll-like receptor 2 | toll like receptor 2 | Plasma Membrane | transmembrane receptor | 7097 | 24088 | 310553 |
| TLR3 | AI957183, CD283, IIAE2, toll-like receptor 3 | toll like receptor 3 | Plasma Membrane | transmembrane receptor | 7098 | 142980 | 364594 |
| TLR7/8 | TLR7 and TLR8 | | Plasma Membrane | group | | | |
| TMPRSS2 | D16Ertd61e, PP9284, PRSS10, transmembrane protease, serine 2, Transmembrane serine protease 2 | transmembrane serine protease 2 | Plasma Membrane | peptidase | 7113 | 50528 | 156435 |
| TNF | AT-TNF, DIF, RATTNF, TMTNF, TNF-a, TNF-alpha, Tnfsf1a, TNFSF2, TNF-α, TNLG1F, tumor necrosis factor, Tumor Necrosis Factor α, tumor necrosis factor, α, tumour necrosis factor, tumour Necrosis Factor Alpha, tumour necrosis factor, alpha, tumour Necrosis Factor α, tumour necrosis factor, α | tumor necrosis factor | Extracellular Space | cytokine | 7124 | 21926 | 24835 |
| Zn2+ | 23713-49-7, zinc 2+, zinc(2+), zinc ion, zinc, ion (Zn2+), Zn+2, Zn(II) | | Other | chemical - endogenous mammalian | PubChem ID: 32051 | | |
